# A genome-wide atlas of recurrent repeat expansions in human cancer

**DOI:** 10.1101/2022.08.24.505159

**Authors:** Graham S. Erwin, Gamze Gürsoy, Rashid Al-Abri, Ashwini Suriyaprakash, Egor Dolzhenko, Kevin Zhu, Christian R. Hoerner, Shannon M. White, Lucia Ramirez, Ananya Vadlakonda, Alekhya Vadlakonda, Konor von Kraut, Julia Park, Charlotte M. Brannon, Daniel A. Sumano, Raushun A. Kirtikar, Alicia A. Erwin, Thomas J. Metzner, Ryan K. C. Yuen, Alice C. Fan, John T. Leppert, Michael A. Eberle, Mark Gerstein, Michael P. Snyder

## Abstract

Expansion of a single repetitive DNA sequence, termed a tandem repeat (TR), is known to cause more than 50 diseases. However, repeat expansions are often not explored beyond neurological and neurodegenerative disorders. In some cancers, mutations accumulate in short tracts of TRs (STRs), a phenomenon termed microsatellite instability (MSI); however larger repeat expansions have not been systematically analyzed in cancer. Here, we identified TR expansions in 2,622 cancer genomes, spanning 29 cancer types. In 7 cancer types, we found 160 recurrent repeat expansions (rREs); most of these (155/160) were subtype specific. We found that rREs were non-uniformly distributed in the genome with an enrichment near candidate cis-regulatory elements, suggesting a role in gene regulation. One rRE located near a regulatory element in the first intron of *UGT2B7* was detected in 34% of renal cell carcinoma samples and was validated by long-read DNA sequencing. Moreover, targeting cells harboring this rRE with a rationally designed, sequence-specific DNA binder led to a dose-dependent decrease in cell proliferation. Overall, our results demonstrate that rREs are an important but unexplored source of genetic variation in human cancers, and we provide a comprehensive catalog for further study.

## Introduction

Expansions of tandem DNA repeats (TRs) are known to cause more than 50 devastating human diseases including Huntington’s disease and Fragile X syndrome^1,2^. TR tracts that cause human disease are typically large (more than 100 base pairs)^1^. However, identifying large TRs with short-read DNA sequencing methods is difficult because they are ubiquitous in the genome, and many are too large—larger than the typical sequencing read length—to uniquely map to the reference genome^3^. Thus, many large TRs go undetected with current genomic technologies, and despite their importance to monogenic disease, the frequency and function of recurrent repeat expansions (rREs) are unknown in complex human genetic diseases, such as cancer^4^.

Previous studies have profiled the landscape of alterations in STRs in cancer genomes^5–7^. In particular, microsatellite instability (MSI)^8–10^, defined by an alteration in the lengths of short TRs, is prevalent in various types of cancer including endometrial (30%), stomach (20%), and colorectal cancers (15%)^5,6,11–12^. However, the systematic analysis of the frequency of genome-wide large TR expansions has not been studied in cancer even though it was posited more than 25 years ago^13^.

Recently, new bioinformatic tools to identify repeat expansions in short-read whole-genome sequencing (WGS) datasets^14–17^ have led to the identification of both known and novel repeat expansions in human disease, primarily in the area of neurological disorders where repeat expansions have historically been studied^14–22^. Here, we analyzed 2,622 human cancer genomes with matching normal samples for the presence of somatic repeat expansions. We identified 160 recurrent repeat expansions (rREs) in seven types of cancer, including many rREs located in or near known regulatory elements. One of these rREs is observed in 34% of kidney cancers, and targeting this repeat expansion with sequence-specific DNA binders led to a dose-dependent decrease in cellular proliferation. Overall, our approach reveals a new class of recurrent changes in cancer genomes and provides an initial resource of these changes. Our results also suggest an opportunity for a new class of oncologic TR-targeted therapeutics^23,24^.

## Results

### Recurrent repeat expansions in cancer

We collected uniformly-processed alignments of whole-genome sequencing (WGS) of tumor-normal pairs in the International Cancer Genome Consortium (ICGC), The Cancer Genome Atlas (TCGA), both a part of the pan-cancer analysis of whole genomes (PCAWG) datasets^25^. After filtering, these data consist of 2,622 cancer genomes from 2509 patients across 29 different cancer types (**Extended Data Figure 1**). Each cancer type was treated as its own cohort and analyzed independent of other cancer types. We called somatic recurrent repeat expansions (rREs) with ExpansionHunter Denovo (EHdn) (see **Methods**), which measures TRs whose length exceeds the sequencing read length in short-read sequencing datasets^26,27^. That is, EHdn performs case-control comparisons using a non-parametric statistical test to determine whether repeat length is longer in tumor genomes compared to matching normal genomes. This approach is analogous to joint population-level genotyping.

We first confirmed the accuracy of EHdn by performing whole-genome short- and long-read sequencing on 786-O and Caki-1 cancer lines. We found that EHdn captured 72% of the repeat expansions observed in long-read sequencing (**Extended Data Figure 2**). We also tested the effect of sequencing coverage on the detection of rREs, and found that EHdn was robust down to 30x coverage (**Extended Data Figure 2**). We then analyzed 2,622 matching tumor and normal genomes with EHdn (285,363 TRs). We identified 578 candidate rREs (locus-level false discovery rate (FDR) < 10%).

EHdn is expected to be sensitive to the copy number variations observed in cancer genomes. We therefore devised and implemented a local read depth filtering method to account for copy number variants, which normalizes the signal originating from repeat reads using the read depth in the vicinity of the TR (see **Methods** and **Extended Data Figure 3**). We benchmarked the local read depth normalization approach with simulated chromosomal amplifications ranging from two (diploid) to 10 copies. We found that this filter accounts for changes in chromosomal copy number in a manner superior to the standard global read depth normalization (**Fig. S5**). Overall, we conclude that local read depth normalization is valuable to identify *bona fide* rREs in cancer genomes and that many of the rREs that pass the filter are expanded in cancer. For example, without local read depth normalization, we could only detect 31% of candidate rREs in independent cohorts of matching tumor-normal tissue samples for breast, prostate, and kidney cancer (15, 18, and 12 patients, respectively). Our local read depth filtering approach removed >75% (418/578) false-positive candidate rREs (**Extended Data Figure 3**). Importantly, several rRE candidates that were removed are situated in hotspots for chromosomal amplification, such as chromosomal 8q amplifications that increase *MYC* production in breast cancer (**Extended Data Figure 3**)^28^. Our analysis suggests that the standalone EHdn method may have selected these loci due to amplification rather than repeat expansions, and thus their removal is important.

After implementing our local read depth filtering strategy, we increased our detection rate to 57% (8/14) in independent cohorts (**Extended Data Figure 3**). Importantly, the loci we could not validate had lower expansion frequencies (5–12%). These rREs may be real but more difficult to validate in the small validation cohorts (**Table S7**). Thus, we believe this number is likely an underestimate of the independent detection rate. Of the 14 candidate rREs that failed our local read depth filter, 29% (4/14) were detected in independent cohorts of samples indicating that the filtering removes most loci that cannot be validated (**Extended Data Figure 3**), but also removes some true positives as well.

After accounting for local read depth, we detected 160 rREs in 7 human cancers (**Fig. 1b**). We expected high concordance with ExpansionHunter given that this tool is related to EHdn, and indeed we observed a 91% confirmation rate with ExpansionHunter (**Extended Data Figure 4**). We found that most (80%) of these loci are rarely expanded in the general population (<5% of the time, *n* = 6,514 genomes, **Extended Data Figure 2**). rREs were primarily observed in prostate and liver cancer, but we also detected rREs in ovarian, pilocytic astrocytoma, renal cell carcinoma (RCC), chromophobe RCC, and squamous cell lung carcinoma. Thus, rREs are found in tissues derived from each of the three primary germ layers (ectoderm, mesoderm, and endoderm), suggesting these expansions are a phenomenon inherent to the human genome rather than any tissue-specific process. In prostate and liver cancer, most cancer genomes (93% and 95%, respectively) contain at least one rRE, with some genomes harboring several rREs (**Fig. 1c**). For some pathogenic repeats, a larger TR length at birth predisposes an individual to somatic repeat expansions later in life^1,2^, but we did not generally observe that with rREs (**Table S8**). Overall, rREs are found in 7 of 29 human cancers examined and are largely cancer subtype-specific.

**Figure 1.**
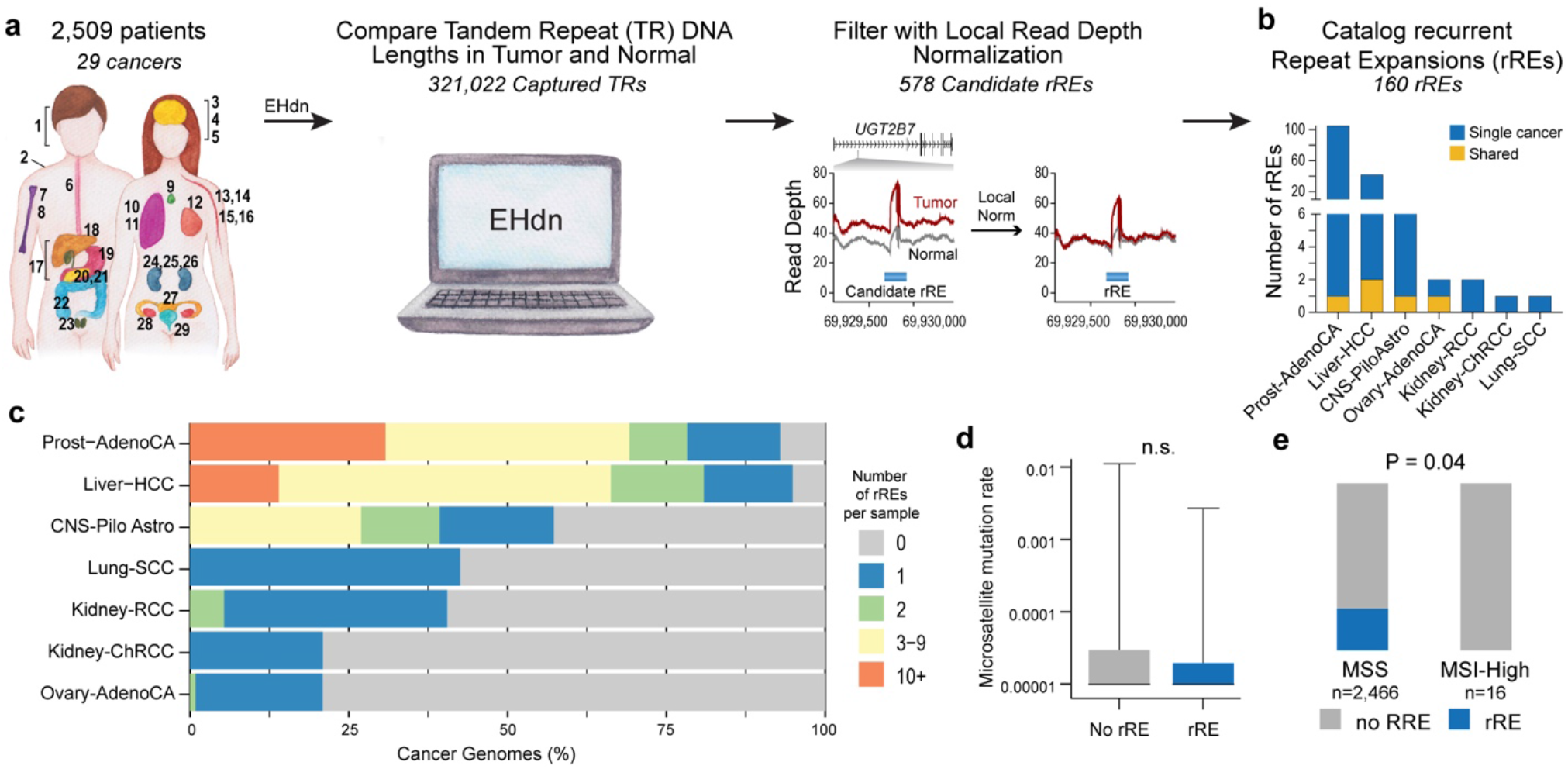
Genome-wide detection of recurrent repeat expansions (rREs) in cancer genomes. a) Scheme of method to identify rREs in 2,509 patients across 29 human cancers. 1, head and neck squamous cell carcinoma (Head−SCC); 2, Skin−Melanoma; 3, glioblastoma (CNS−GBM); 4 medulloblastoma (CNS−Medullo); 5, pilocytic astrocytoma (CNS−PiloAstro); 6, esophageal adenocarcinoma (Eso−AdenoCA), 7, osteosarcoma (Bone−Osteosarc); 8, leiomyosarcoma (Bone−Leiomyo); 9, thyroid adenocarcinoma (Thy−AdenoCA); 10, lung adenocarcinoma (Lung−AdenoCA); 11, lung squamous cell carcinoma (Lung−SCC); 12, mammary gland adenocarcinoma (Breast−AdenoCA); 13, B-cell non-Hodgkin lymphoma (Lymph−BNHL); 14, chronic lymphocytic leukemia (Lymph−CLL); 15, acute myeloid leukemia (Myeloid−AML); 16, myeloproliferative neoplasm (Myeloid−MPN); 17, biliary adenocarcinoma (Biliary-AdenoCA); 18, hepatocellular carcinoma (Liver−HCC); 19, stomach adenocarcinoma (Stomach−AdenoCA); 20, pancreatic adenocarcinoma (Panc−AdenoCA), 21, pancreatic neuroendocrine tumor (Panc−Endocrine); 22, colorectal adenocarcinoma (ColoRect−AdenoCA); 23, prostatic adenocarcinoma (Prost−AdenoCA); 24, chromophobe renal cell carcinoma (Kidney−ChRCC); 25, renal cell carcinoma (Kidney−RCC); 26, papillary renal cell carcinoma (Kidney−pRCC); 27, uterine adenocarcinoma (Uterus−AdenoCA); 28, ovarian adenocarcinoma (Ovary−AdenoCA); 29, transitional cell carcinoma of the bladder (Bladder−TCC). b) Distribution of rREs across cancer types. c) Proportion of cancer genomes with rREs. d) STR mutation rate for cancer genomes with and without a rRE. Two-tailed Wilcoxon rank sum test. e) Distribution of rREs across microsatellite stable (MSS) and microsatellite instability high (MSI-high) cancers. Chi-square test with Yates’ correction.

We next examined whether rREs correlate with changes in MSI^5,6^. We determined whether samples harboring an rRE had a higher mutation rate in STRs, which is a hallmark of MSI^5,29^. We did not observe any significant difference in STR mutation rate for genomes with an rRE compared to those lacking an rRE (two-tailed Wilcoxon rank sum test, *P* = 0.27, **Fig. 1d**). We also compared cancer genomes harboring rREs with cancer genomes previously identified as MSI, using recent results from the PCAWG consortium^29^. We did not observe any enrichment in MSI for samples harboring an rRE, and instead found a weak but significant preference for rREs in microsatellite stable (MSS) samples, not MSI samples (Two-tailed Wilcoxon rank-sum test, *P* = 0.04, **Fig. 1e**, see also **Extended Data Figure 5**). Thus, our findings might suggest a model where rREs are formed by a process that is distinct from MSI.

In addition to MSI, different mutational processes lead to a signature of somatic mutations. We tested whether rREs are associated with known mutational signatures by comparing them to 49 signatures of single base substitutions (SBS) and 11 doublet base substitutions (DBS)^30^. We performed a multiple linear regression to predict the number of rREs in a sample based on SBS and DBS signatures, respectively. Only one DBS signature, DBS2, showed a very weak association with rREs (r^2^ = 0.12) (**Extended Data Figure 5**).

### rREs overlap regulatory elements

Among the 160 rREs, we observed a variety of different motifs (**Table S1**) whose repeat unit length follows a bimodal distribution, consistent with REs identified in other diseases (**Fig. 2a, Extended Data Figures 6 and 7**)^27^. rREs are distributed across a range of GC content; approximately half (76/160) have GC content less than 50% (**Table S1**). Six rREs contained a known pathogenic motif, all of which were GAA^31^. We examined if any motifs were enriched in the rRE catalog compared to the tandem repeat finder (TRF) catalog. Although this enrichment could arise from a biological and/or technical process, we found that one of the three enriched motifs was GAA (**Fig. 2b**). As an example, Friedreich’s ataxia is caused by a repeat expansion of a GAA motif in the intron of the frataxin gene. This expansion leads to DNA methylation and the deposition of repressive chromatin marks, leading to robust repression of the gene and development of disease^31^. Because of this, we suspect some of the rREs found in cancer might alter the epigenome and affect gene regulatory networks.

**Figure 2.**
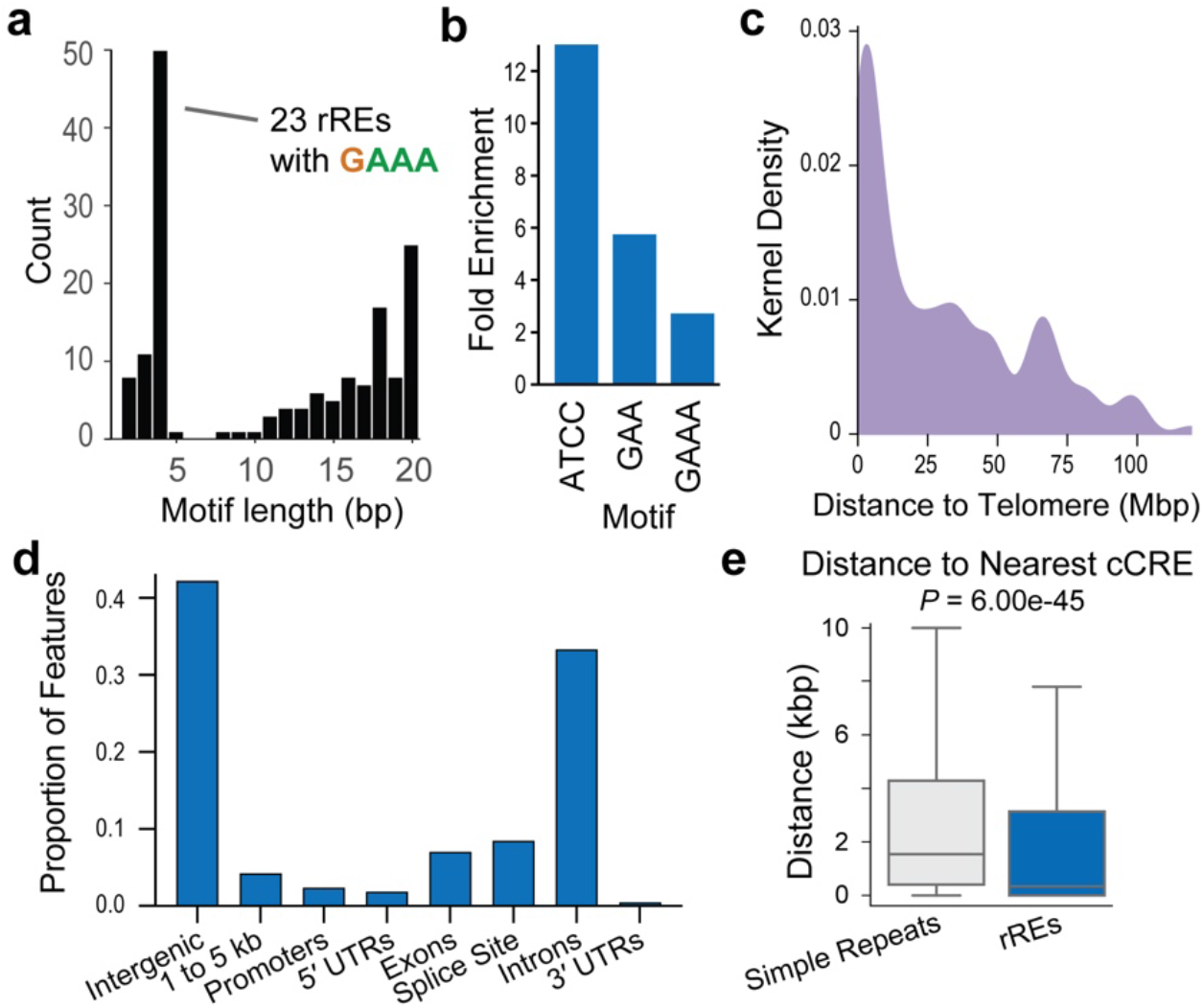
Features of rREs. a) Distribution of the repeat unit (motif) for rREs. b) Motifs enriched in the catalog of rREs. c) Distance of rREs to the end of the chromosome arm. d) Proportion of genic features that overlap with rREs. UTR, untranslated region. e) Distance of simple repeats and rREs to the nearest ENCODE candidate cis-regulatory element (cCRE). Center values represent the median. Welch’s *t*-test.

rREs were distributed non-uniformly across the genome, with a bias towards the ends of chromosome arms (**Fig. 2c, Extended Data Figure 6**). This observation is consistent with previous reports of TRs and structural variants^15,32^. We also examined the distribution of rREs relative to gene features with annotatr (**Fig. 2d**)^33^. The 7% of rREs labeled as exonic appeared proximal to, but not within, exons, but others were in introns, untranslated regions (UTRs), and splice sites. These results suggest rREs may play different functional roles in the regulation of gene expression.

We measured the distance between rREs and candidate cis-regulatory elements (cCREs)^34^; cCREs comprise approximately one million functional elements including promoters, enhancers, DNase-accessible regions, and insulators bound by CCCTC-binding factor (CTCF). An rRE near a regulatory element could alter the function of that regulatory element, as is observed in Fragile X syndrome and Friedreich’s ataxia^1^. Interestingly, rREs are located closer to cCREs than expected by chance, and we find that 47 of 160 rREs directly overlap with a known cCRE (Welch’s *t-*test, *P* = 6.00e-45, **Fig. 2e & Extended Data Figure 7**). Thus, rREs are often found in or near functional regions of the genome.

### rREs with links to cancer

We mapped each rRE to the nearest genes and found that nine rREs map to Tier 1 genes present in the census of somatic mutations in cancer (COSMIC) database (**Fig. 3, Table S1**). We also observed a strong correlation with cancer-related genes (Jensen disease-gene associations^35^). That is, four of the top five diseases associated with the collection of 160 rRE are cancers (**Fig. 3b, Table S4**).

**Figure 3.**
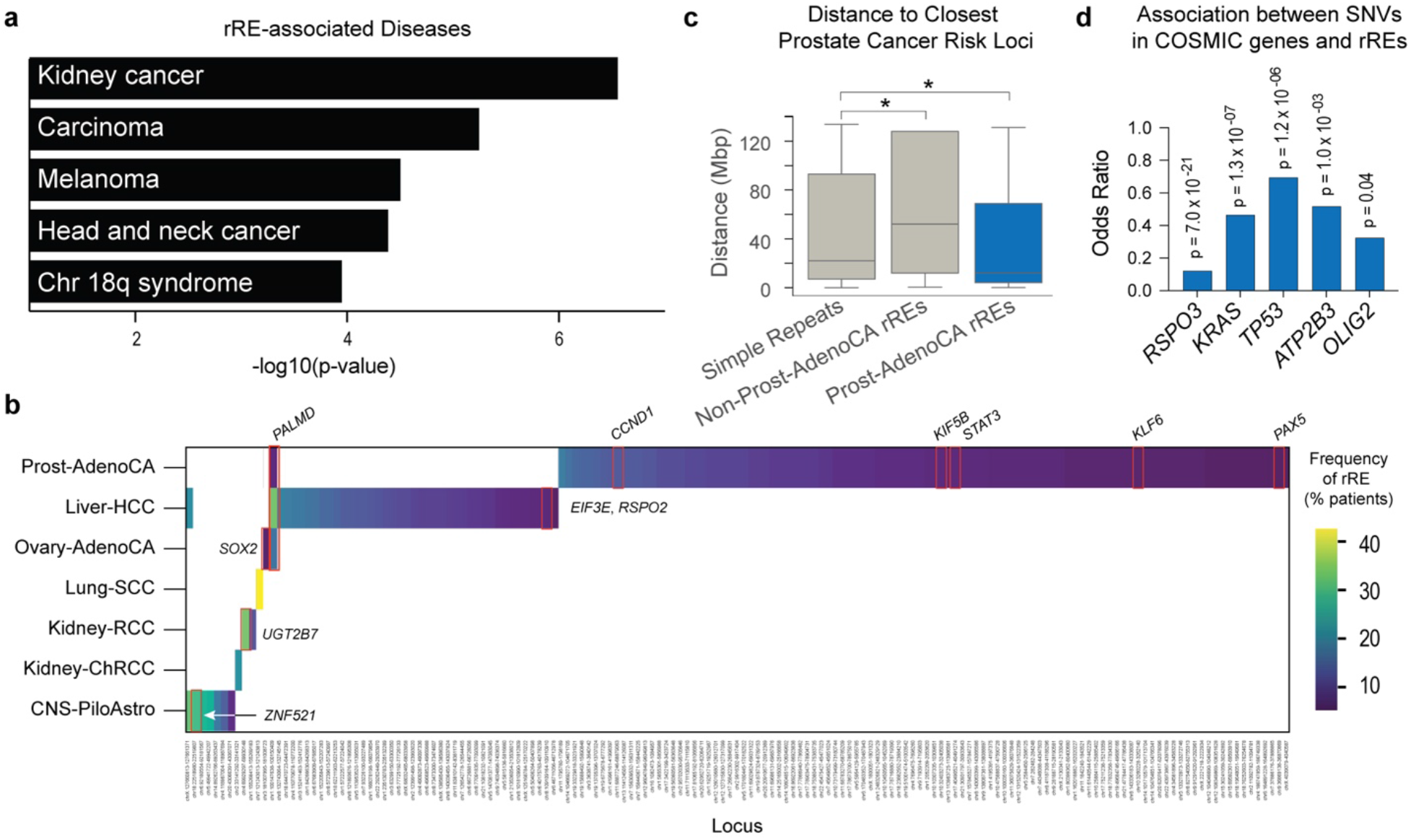
Association of rREs with cancer features. a) Association of rREs with human diseases. b) Frequency of rREs in genes of interest, including the nine COSMIC genes, are highlighted. c) Distance of simple repeats, non-prostate cancer rREs, and prostate-cancer rREs to the nearest prostate cancer risk locus. Center values represent the median. Statistical significance was measured with Welch’s *t*-test (* q < 0.10). d) Association between SNVs in genes in the census of somatic mutations in cancer (COSMIC) Tier 1 genes and the presence of rREs. Student’s *t*-test with FDR correction by Benjamini-Hochberg.

To examine whether some rREs play a role in oncogenesis, we looked at their association with previously-identified cancer risk loci. Many rREs were identified in prostate cancer, and 63 loci have previously been associated with susceptibility to prostate cancer from available genome-wide association studies^36^. When we examined the co-localization of rREs and cancer risk loci in prostate cancer, we found that rREs are located closer to prostate cancer susceptibility loci than standard STRs or by chance (Student’s *t*-test, FDR q = 0.08, **Fig. 3c & Extended Data Figure 7**).

We next studied the relationship between the occurrence of the census of somatic mutations in cancer (COSMIC) genes to the occurrence of rREs (**Fig. 3d**). Interestingly, we found that there are five COSMIC genes whose somatic mutations are found to occur significantly more in patients’ genomes with no rREs, after correcting for multiple hypothesis testing. Among them, *TP53* was particularly striking, as wildtype *TP53* is critical for mediating the pathogenic effects of repeat expansions in both Amyotrophic Lateral Sclerosis (ALS) and Huntington’s Disease^37,38^. Consistent with these findings, a DNA damage repair gene in yeast, *Rad53*, is phosphorylated and activated in the presence of an expanded repeat^39^.

MSI-high cancers are often correlated with higher levels of immune cell infiltration^40^. We hypothesized that some rREs might also be associated with higher immune cell infiltration, but we did not observe a correlation between cytotoxic activity^41^ and the presence of an rRE (**Extended Data Figure 9**). Because there are matching RNA-seq data for only 4 of 160 rREs, this analysis warrants further investigation as more matching WGS and RNA-seq datasets become available.

### An rRE in the *UGT2B7* gene observed in RCC

A GAAA expansion located in the intron of *UGT2B7* was observed in 34% of RCC samples. *UGT2B7* is a glucuronidase that clears small molecules—including chemotherapeutics—from the body and is selectively expressed in the kidney and liver^42^.

With gel electrophoresis, we identified the expected TR size of ∼26 GAAA repeats in the normal kidney cell line, HK-2, corresponding closely to the length observed in the reference genome (**Fig. 4a**). In contrast, we identified an expansion between ∼63 and ∼160 GAAA repeat units in 5 of 8 clear cell RCC cell lines. Most expansions were heterozygous (**Fig. 4a**). Long-read DNA sequencing with highly-accurate PacBio HiFi reads confirmed the PCR results and revealed the precise structure of this repeat expansion at single base pair resolution for both 786-O and Caki-1 (**Fig. 4b**). We also detected this repeat expansion in five out of 12 primary kidney tumor tissue samples from patients with clear cell RCC (**Extended Data Figure 8**), which showed more heterogeneity than the RCC cell lines; more heterogeneity might be expected for human tumor samples compared to the clonal cell lines.

**Figure 4.**
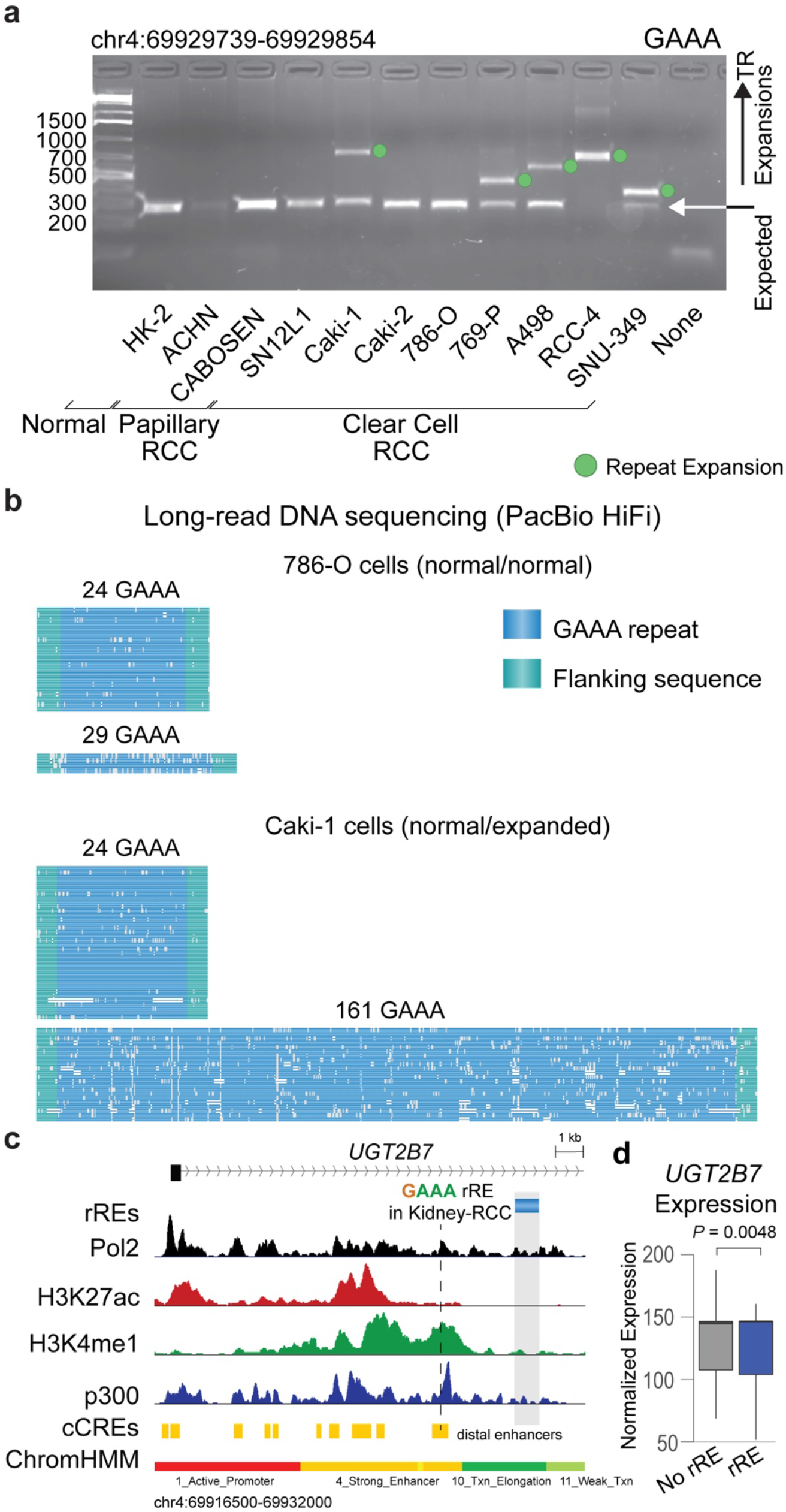
An rRE in Renal Cell Carcinoma (RCC). a) Gel electrophoresis of the GAAA tandem repeat in RCC samples. This analysis was performed in duplicate and the gel is representative of the results. For gel source data, see **Fig. S1**. b) Visualization of long-read sequencing of GAAA rRE in the intron of *UGT2B7*. Data are from PacBio HiFi sequencing. c) The locus surrounding the rRE detected in the intron of *UGT2B7*. Signal traces of Pol2, H3K27ac, H3K4me1, and p300 in HepG2 cells are shown. Candidate cis-regulatory elements (cCREs) and chromatin states (ChromHMM) are also depicted. d) Expression of *UGT2B7* isoform ENST00000508661.1 in RCC samples as a function of the detection of the rRE in *UGT2B7* (Normalized Expression, Counts). Center values represent the median. Significance was measured by Wald test with FDR correction (Benjamini-Hochberg).

Given that *UGT2B7* is selectively expressed in the liver and kidney, and that it plays a role in clearing small molecules from the body, we examined whether this rRE may be located near any functional elements that could regulate its expression. Analysis of the chromatin environment surrounding the rRE in *UGT2B7* revealed a nearby enhancer, raising the possibility that this rRE alters the expression of *UGT2B7* (**Fig. 4c**). The repeat motif of this rRE, GAAA, appears similar to the pathogenic repeat motif found in Friedreich’s ataxia, which is GAA. The pathogenic GAA repeat expansion blocks *FXN* expression^31^. We therefore hypothesized that the intronic GAAA repeat expansion might repress the expression of *UGT2B;* we found a modest decrease in expression that was not statistically significant (**Extended Data Figure 8**). While this rRE is also not associated with a difference in survival (**Extended Data Figure 8**), it is associated with a significant decrease in a transcript isoform in *UGT2B7* (Wald test with FDR correction, *P* = 0.0048) (**Fig. 4e**). Interestingly, a shift in isoform usage of *UGT2B7* has been noted in cancer^43^.

### Repeat-targeting anti-proliferative agents

Do GAAA repeat expansions contribute to cell proliferation? We previously showed that targeting a related TR motif, GAA, with synthetic transcription elongation factors (Syn-TEF1) reverses pathogenesis in several models of Friedreich’s ataxia^23^. Therefore, if the GAAA rRE in RCC behaves similarly, then a Syn-TEF targeting GAAA may have anti-proliferative activity. We rationally designed Syn-TEF3, which contains a GAAA-targeting polyamide (PA), and a bromodomain ligand, JQ1, designed to recruit part of the transcriptional machinery (**Fig. 5a** and **Fig. S2**). We also included a control molecule, Syn-TEF4, which targets G**<u>G</u>**AA TRs, as well as polyamides (PAs) PA3 and PA4 that lack the JQ1 domain. We have previously shown that Syn-TEFs and PAs localize to repetitive TRs in living cells^23,44^.

**Figure 5.**
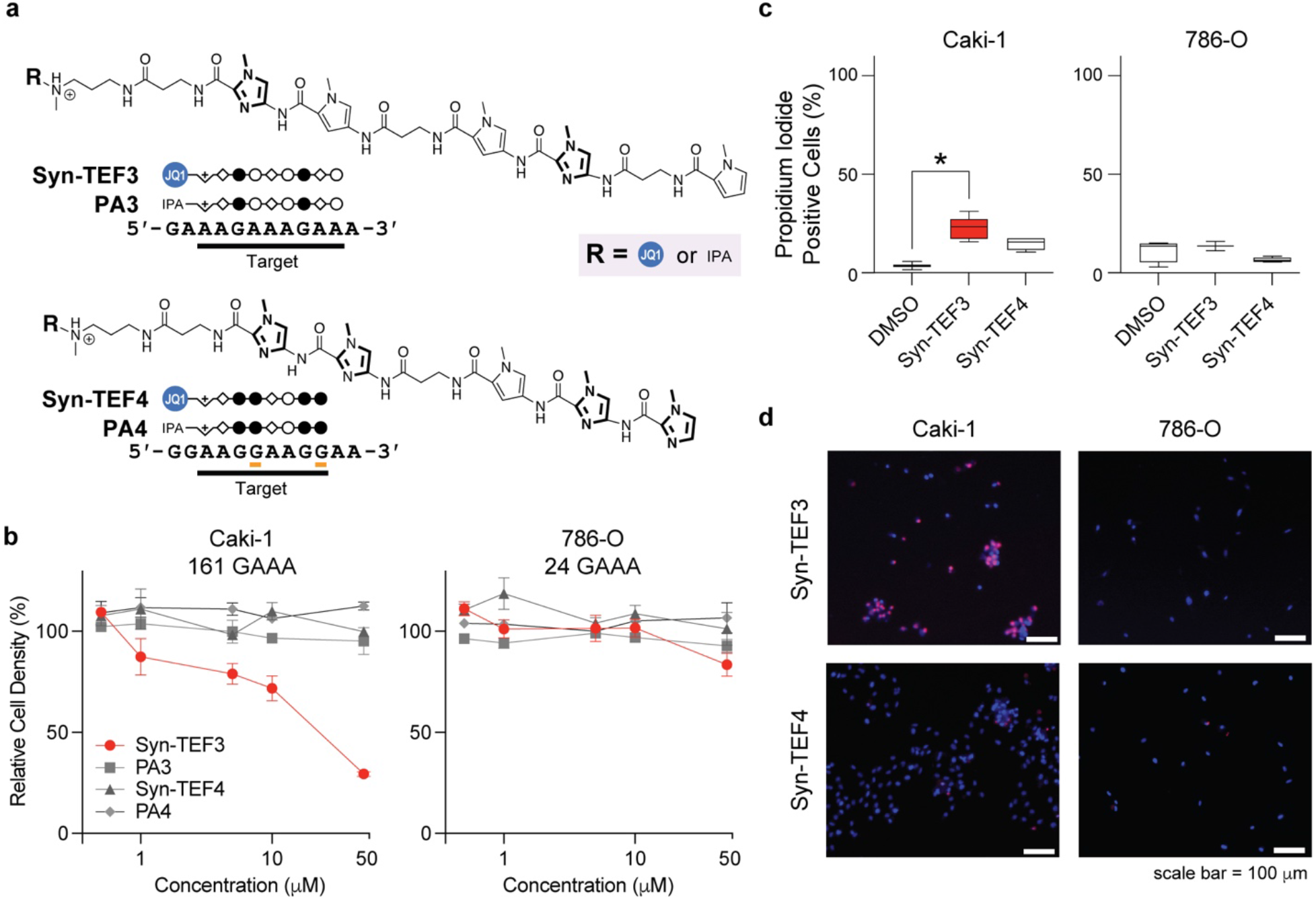
The design and characterization of GAAA-targeting molecules in RCC. a) Chemical structures of Syn-TEF3, PA3, Syn-TEF4 and PA4. Syn-TEF3, and PA3 target 5′- AAGAAAGAA-3′. Syn-TEF4* and PA4 target 5′-AAGGAAGG-3′. The structures of *N*- methylpyrrole (open circles), *N*-methylimidazole (filled circles), and β-alanine (diamonds) are shown. N-methylimidazole is bolded for clarity. The structure of JQ1 linked to polyethylene glycol (PEG_6_) is represented as a blue circle. The structure of isophthalic acid and linker is represented as IPA. Complete chemical structures are depicted in **Fig. S2**. The asterisk indicates the site where the R group attaches to the polyamide. Mismatches formed with Syn-TEF4 and PA4 are indicated with orange. b) Relative cell density of RCC cell lines Caki-1 and 786-O following treatment (72 h) with compounds, as indicated. Relative cell density was measured with CCK-8 assay (see **Methods**). Results are mean ± SEM (*n* = 4). c) Quantitation of the percentage of propidium iodide-positive cells. Whiskers are minimum and maximum values. * p < 0.05. P values are from a one-way ANOVA with multiple comparisons. d) Live cell microscopy of Caki-1 and 786-O cells stained with propidium iodide (red) and Hoechst 33342 (blue). Scale bars, 100 µm. See also **Extended Data Figure 10**.

The effect of Syn-TEFs on cell proliferation was examined (**Fig. 5b**). Caki-1 and 786-O were selected because they have the largest (161) and smallest (24) GAAA tracts within the first intron of *UGT2B7*, respectively. In a dose-dependent manner, we observed that Syn-TEF3 led to a significant decrease in the proliferation of Caki-1 cells but had little effect on 786-O cells. Syn-TEF4, which does not target a GAAA TR, did not significantly decrease proliferation in either of the cell lines tested, demonstrating the requirement for GAAA-specific targeting (**Fig. 5b**). Two additional GAAA repeat expansion cell lines as well as two additional control non-expanded lines showed a similar association between Syn-TEF sensitivity and the presence of the repeat expansion (**Extended Data Figure 10**). Consistent with this finding, Caki-1 cells treated with Syn-TEF3 exhibited a significant increase in cell death compared to DMSO control, as measured by propidium iodide staining (**Fig. 5c,d** and **Extended Data Figure 10**). In contrast, 786-O cells treated with Syn-TEF3 showed no significant difference in propidium iodide-positive cells compared to DMSO (**Fig. 5c,d** and **Extended Data Figure 10**). Importantly, the Syn-TEF4, PA3, and PA4 control agents exhibited no significant effect on cell death in either cell line compared to vehicle control (**Fig. 5c,d** and **Extended Data Figure 10**). These results suggest that GAAA repeat expansions may represent a genetic vulnerability in RCC and provide a proof-of-principle study for the functional role of rREs in cancer.

## Discussion

Here, for the first time, we conduct a genome-wide survey of recurrent repeat expansions (rREs) across cancer genomes, distinct from MSI. Our data identified (i) 160 rREs in 7 human cancers and revealed that (ii) most (155 of 160) rREs are cancer subtype-specific; (iii) amongst diseases, rREs are enriched in human cancer loci and tended to occur near regulatory elements; (iv) recurrent repeat expansions do not correlate with MSI status; and (v) targeting a GAAA repeat expansion in RCC with a small molecule leads to cancer cell killing. Taken together, our results uncover an unexplored genetic alteration in cancer genomes with important mechanistic and therapeutic implications.

Cancer cells evolve and adapt in response to environmental or pharmacological perturbations, but the mechanisms supporting these changes are still being uncovered. One source of genetic variation that may enable genetic adaptations is TR DNA sequences. Mutations in repeat length of TRs can occur up to 10,000 times more frequently than single nucleotide variants (SNVs) or insertions and deletions (INDELs)^1^. Repeat expansions may provide a source of genetic variation to enable cancer cells to adapt to changes in the environment^45^. Indeed, colorectal cancers acquire mutations in STRs in response to targeted therapy just 24 hours following treatment, suggesting that mutations in these regions may associate with rapid evolution^46^. In future studies, it will be particularly valuable to study repeat expansions in the genomes of cancer cells that face changing environments, including metastasis and chemotherapy.

Historically, MSI has been the focus of efforts to profile changes in STRs in cancer genomes because specific cancer-causing genetic alterations in repair genes can promote widespread STR alterations. Interestingly, we find little to no correlation between rREs and MSI. These results are consistent with previous findings in which the correlation between MSI and repeat instability at larger TRs is not definitive^47^. MSI may contribute to a subtype of rREs that we have not yet uncovered, or rREs may arise from a mutation process that is distinct from that of MSI. There are several different DNA cellular repair systems, and presumably the rREs that we observed are due to very specific loci-associated mechanisms or activities. Some of these repeat expansions may be due to cis-regions with interesting DNA or chromatin configurations that are prone to expansion at distinct loci, rather than gene mutations that cause global trans effects, as occurs in MSI.

There are numerous mechanisms by which a repeat expansion can alter cellular function. Known pathogenic repeat expansions can alter the coding sequence of a protein, such as in the case of Huntington’s disease. However, there are several repeat expansions that occur in non-coding regions that alter gene expression^1^. In other instances, the repeat expansion can lead to a pathogenic RNA molecule (myotonic dystrophy) or protein (ALS)^1^. Finally, repeat expansions in MSI-associated cancers, which are too small to detect by EHdn, can disrupt DNA replication^48^. Thus, our catalog represents a powerful resource to explore the mechanisms by which rREs alter cellular function in cancer.

The identification of repeat expansions would benefit from improved sequencing coverage and increased cohort sizes. Like other tools that identify repeat expansions, we cannot distinguish zygosity from sample heterogeneity or obtain precise lengths of repeats. Our independent experimental validation showed that some repeat expansions are heterogeneous (**Extended Data Figure 8**). We suspect that tumor heterogeneity may lead to an underreporting of rREs. Furthermore, this study focuses on somatic mutations, but repeat expansions that occur in the context of normal development will be another important area of study^4^. Furthermore, germline events that predispose an individual to cancer would also be worthwhile to study; there is evidence that a TR in the androgen receptor gene is associated with prostate cancer onset, tumor stage, and tumor grade^49^. Finally, we only detected changes in repeat length that were greater than sequencing read length. In future studies, it will be important to explore recurrent changes that are smaller in length. Finally, it is important to acknowledge that rREs could be mediators of phenotypes or passengers that result from genetic instability and clonal selection. In the one instance where we targeted the rRE in RCC, cell proliferation was reduced, consistent with a mediator role for this rRE. Distinguishing between these two possibilities for each rRE is an important line of work in the future.

To our knowledge, this is the first genome-wide survey of repeat expansions beyond a neurological or neurodegenerative disorder. Thousands of high-quality whole genome sequences exist for many diseases, and our data provide evidence that repeat expansions should be explored beyond the classical bounds of neurodegenerative diseases where they have been most investigated.

A careful dissection of repeat expansions in human disease may reveal their role as causative or contributory. We show here that repeat expansions can be targeted by tandem repeat-targeting precision molecules^23^. Thus, our results set the stage for a new class of therapeutics to be deployed in cancer and other diseases.

## Methods

### Data curation

We obtained white-listed data from the International Cancer Genome Consortium (ICGC) and The Cancer Genome Atlas (TCGA) pan-cancer analysis of whole genomes (PCAWG) dataset. Data were accessed through the Cancer Genome Collaboratory. We used the aligned reads (BAM files), which were aligned to GRCh37 as described previously^25^. These data are available through the PCAWG data portal (https://docs.icgc.org/pcawg). A list of samples included in the analysis is available in **Table S2**.

### Identification of somatic recurrent repeat expansions

We analyzed tumor and matching normal samples for each cancer type independently. We executed ExpansionHunter Denovo (EHdn) (v0.9.0)^15^ with the following parameters: --min-anchor-mapq 50 --max-irr-mapq 40. To prioritize loci, we developed a workflow termed Tandem Repeat Locus Prioritization in Cancer (TROPIC). We included loci from chr1-22, X, and Y for downstream analysis. We removed loci where >10% of Anchored in-repeat read (IRR) values were >40, which is the theoretical maximum value. The p-value (a non-parametric one-sided Wilcoxon rank sum test) for each locus was used to calculate a false discovery rate (FDR) q-value. Loci with FDR < 0.10 are reported. We selected loci where >5% of samples had an Anchored in-repeat read (IRR) Quotient > 2.5. For a repeat expansion to be detected by EHdn, the tandem repeat must be larger than the sequencing read length. A somatic repeat expansion was defined as having an FDR q-value < 0.05 between tumor and normal samples. To call repeat expansions in individual cancer samples, we analyzed the distribution of tumor and normal Anchored IRR values and selected a conservative threshold for the Anchored IRR Quotient ((Tumor Anchored IRR – Normal Anchored IRR)/(Normal Anchored IRR + 1)) > 2.5 (**Extended Data Figure 4**).

### Local read depth normalization

EHdn normalizes the number of Anchored IRRs for a given locus to the global read depth. To account for chromosomal amplifications and other forms of genetic variation that could alter local read depth, we performed the following normalization. For each rRE locus and sample in its corresponding cancer, samtools v1.13 was used with the parameter depth -r to find the read depth at each base pair within the locus and a 500 bp region surrounding the start and stop positions of the TR. We calculated the average read depth at each base pair and defined this as the local read depth. Finally, we calculated the local read depth-normalized Anchored IRR value specific to a sample and rRE combination by dividing the unnormalized Anchored IRR value from EHdn by the local read depth at the locus.

### Generation of CABOSEN cells

CABOSEN cells were generated from a <u>cabo</u>zantinib-<u>sen</u>sitive (CABOSEN) human papillary RCC xenograft tumor grown in RAG2^-/-^ gammaC^-/-^ mice, as described previously^50^. Tumor tissue was minced with a sterile blade and the cell suspension cultured in DMEM/F-12 medium (Corning) supplemented with 10%(v/v) Cosmic Calf Serum (ThermoFisher). Cells were expanded and cryopreserved in growth medium supplemented with 10%(v/v) DMSO and cells from passage 8 were used for analysis.

### Analysis of rREs by gel electrophoresis

We performed PCR with CloneAmp HiFi PCR Mix (Takara Biosciences, Mountain View, CA) and added DMSO to a final concentration of 5-10% (v/v) as needed. All cell lines were tested negative for mycoplasma contamination with the MycoAlert Mycoplasma Detection Kit (Lonza). Cell line identities were authenticated by STR profiling by the Genetic Resources Core Facility at Johns Hopkins University, with the exception of SNU-349, which did not match the reported STR profile of SNU-349 or any other catalogued cell line, but has a mutated *VHL* gene and expresses high levels of *PAX8* and *CA9*, consistent with ccRCC origin. A list of primers used to analyze the loci is available in **Table S6**.

### Visualization of repeat expansions with ExpansionHunter and REViewer

To inspect the reads supporting a repeat expansion, we annotated the repeat as described on the GitHub page for ExpansionHunter. We then profiled the region with ExpansionHunter (v4.0.2) using the default settings^14^. The resulting reads were visualized with REViewer (v0.1.1) using the default settings. REViewer is available at https://github.com/Illumina/REViewer. A repeat expansion was called when the repeat tract length for one allele of the tumor sample was greater than 100 bp and exceeded the repeat tract length of either normal allele. A locus was considered validated if at least 10 cancer genomes had a repeat expansion.

### Validation of rREs in independent cohorts of samples

Twelve pairs of matching normal and tumor samples from patients with clear cell renal cell carcinoma were obtained with the patients’ informed consent *ex vivo* upon surgical tumor resection (Stanford IRB-approved protocols #26213 and #12597) and analyzed. Eighteen and 15 pairs of matching normal and tumor samples for prostate and breast cancer, respectively, were obtained from the Tissue Procurement Shared Resource facility at the Stanford Cancer Institute and analyzed. Nucleic acid was isolated with either the Quick Microprep Plus kit (Catalog D7005) or the Zymo Quick Miniprep Plus kit (Catalog D7003) (Zymo Research, Irvine, CA). Gel electrophoresis was performed as described above. A locus was considered detected if a somatic repeat expansion was identified in at least one patient tumor sample compared to a matching normal sample.

### Downsampling Analysis

For the downsampling analysis, tumor genomes from renal cell carcinoma samples were downsampled from their mean (52x) sequencing depth to 40, 30, 20, and 10x with the samtools view command. EHdn was run, as described above for each of the sequencing depths, and the Bonferroni-corrected p-value was plotted for the recurrent repeat expansion in *UGT2B7* (GAAA, chr4:69929297-69930148).

### Benchmarking the Local Read-Depth Normalization (LRDN) filter

We benchmarked the local read depth filter *in silico* by observing its behavior with simulated reads. First, we created a reference genome containing artificially expanded repeats. We randomly selected 10 TRs located in chromosome 1 that were less than the sequencing read length of 100 bp. We artificially expanded these TRs in chromosome 1 of GRCh37 with the BioPython python package (version 1.79). Next, we used wgsim (version 0.3.1-r13) to simulate reads from the reference file with the command “wgsim -N 291269925 -1 100 -2 100 reference_file.fasta output.read1.fastq output.read2.fastq”. The number of reads (specified by the -N option) was calculated to achieve 30x coverage of chromosome 1. The resulting pair of files, hereinafter referred to as the base fastq files, contained a copy number of 2 for all of the expansions.

To simulate copy number amplification, the read simulation process was repeated using reference files that contained only the artificially expanded repeats and their surrounding 1,000 bp flanks. We created 10 pairs of fastq files, each with an increasing copy number. We specified the copy number by multiplying the number of reads to generate (wgsim -N option) by the required number. To generate the final set of fastq files, we concatenated each pair of copy number-amplified fastq files with the base fastq files. The end result is 8 pairs of fastq files that contain reads of chromosome 1 and a copy number amplification varying from 2 to 10 of the expanded repeats.

The base fastq file with a copy number of 2, in addition to the eight copy number-amplified fastq files, were aligned to chromosome 1 of GRCh37 with bwa-mem (v 0.6) with the default options. The resulting SAM files were converted to BAM format with samtools (v 1.15) with the default options. Finally, we ran the EHdn profile command (v 0.9.0) with the minimum anchor mapping quality set to 50 and maximum IRR mapping quality set to 40. Finally, the Anchored IRR values were extracted by overlapping the STR coordinates with the *de novo* repeat expansion calls.

### Short-read and long-read DNA sequencing

We sequenced Caki-1 and 786-O with both short-read sequencing (60x sequencing coverage, 150 bp paired-end sequencing on a NovaSeq 6000 instrument) and long-read DNA sequencing (50x sequencing coverage, PacBio HiFi sequencing on a Sequel IIe instrument). We aligned the long reads to GRCh37 with pbmm2 v1.7.0, using the parameters --sort --min- concordance-perc 70.0 --min-length 50. We aligned the short reads to GRCh37 with Sentieon (v202112.01) with parameters -K 10000000 -M, an implementation of BWA-MEM, and analyzed the samples with EHdn, as described above. We included loci containing at least one sample with an Anchored IRR value >0 for further analysis. Anchored IRR values >0 arise when the repeat length exceeds the sequencing read length. To benchmark EHdn against long-read sequencing data, we manually determined the TR length of a given locus in the long-read sequencing data. If the TR length in the long-read sequencing data exceeded the short-read sequencing read length of 150 bp, we considered that locus confirmed.

The PacBio HiFi data were aligned to GRCh37 with pbmm2 v1.7.0 and visualized at the *UGT2B7* locus with Tandem Repeat Genotyper v0.2.0 (https://github.com/PacificBiosciences/trgt).

### Analysis of rRE loci

To determine if rREs were associated with any human diseases, rREs were mapped to genes with GREAT (v4.0.4, default settings)^51^. The resulting genes were analyzed with Enrichr using Jensen Diseases^52^. To determine whether repeat expansions were associated with microsatellite instability-high (MSI-High) cancers, we obtained data from Hause et al^5^. The percentage of MSI-high cancers was obtained from colon adenocarcinoma (COAD), stomach adenocarcinoma (STAD), kidney renal cell carcinoma (KIRC), ovarian serous cystadenocarcinoma (OV), prostate adenocarcinoma (PRAD), head and neck squamous cell carcinoma (HNSC), liver hepatocellular carcinoma (LIHC), bladder urothelial carcinoma (BLCA), glioblastoma multiforme (GBM), skin cutaneous melanoma (SKCM), thyroid carcinoma (THCA), and breast invasive carcinoma (BRCA) and compared to the number of repeat expansions and the percentage of patients with at least one repeat expansion in the corresponding cancer type from the PCAWG dataset. We also overlapped cancer genomes containing rREs with the microsatellite mutation rate, which we term the STR mutation rate, and MSI calls from Fujimoto et al^29^. The association of rREs with STR mutation rate was assessed with the two-tailed Wilcoxon rank sum test. The association of rREs with MSI calls was assessed with Chi-square test with Yates’ correction.

To determine whether rREs are associated with known mutational signatures, we downloaded mutational signatures from the ICGC DCC (https://dcc.icgc.org/releases/PCAWG/mutational_signatures/Signatures_in_Samples). We performed a multiple linear regression for each single-base-substitution (SBS) and doublet-base-substitution (DBS) signatures to identify predictors of the number of rREs present in a sample. To choose the predictors, we performed best subset selection on DBS and SBS signatures and included age as a possible confounding factor. We used the statsmodels v0.12.2 in Python and, specifically, the ordinary least squares model found in the statsmodels.api.OLS module to estimate the coefficients of the selected predictors in their corresponding multiple linear regression model^53^.

To determine whether repeat expansions were associated with a difference in cytotoxic activity, we calculated cytotoxic activity as previously described for four cancers that had matching RNA-seq and WGS^41^. For each locus, we compared cytolytic activity for patients with a repeat expansion to patients without a detected repeat expansion using a Welch’s *t*-test with correction for multiple hypothesis testing (Benjamini-Hochberg FDR q-value < 0.05). rREs were annotated with genic elements using annotatr (v1.18.1)^33^.

To determine if rREs were associated with regulatory elements, we downloaded candidate cis-regulatory elements (cCREs)^34^ and mapped them to GRCh37 with LiftOver (UCSC)^54^. We determined the distance between rREs and cCREs with the bedtools closest command (v2.27.1)^55^, and compared this distance to the simple repeats catalog^56^. To compare the distance to ENCODE cCREs, a Welch’s *t*-test was performed.

To determine if prostate cancer rREs were associated with prostate cancer susceptibility loci^36^, we calculated the distance to three sets of loci using the “bedtools closest” command. We calculated the distance between (1) rREs present in prostate cancer samples and prostate cancer susceptibility loci, (2) rREs not present in cancer samples and cancer susceptibility loci, and (3) simple repeats and cancer susceptibility loci. To compare the distances between these three associations, we performed a Welch’s *t*-test with FDR correction (Benjamini-Hochberg).

To determine whether rREs were associated with replication timing, we downloaded Repli-seq replication timing data from seven cell lines from the ENCODE website (NCI-H460, T470, A549, Caki2, G401, LNCaP, and SKNMC)^57^. We selected regions for which all cell lines had concordant signals for analysis (early or late replication designations agreed for each cell line at a given locus). We determined whether there was a difference in the distribution of rREs across early- and late-replicating regions compared to the simple repeats catalog with bootstrapping (*n=*10,000). We sampled 54 loci (the number of rREs that are present in a concordant replication region) from rREs and simple repeats. A Welch’s t-test was performed on the bootstrapped samples to estimate a *p*-value. We applied FDR correction (Benjamini-Hochberg) to the estimated p-values. To determine whether rRE status in *UGT2B7* was associated with survival outcome in clear cell RCC patients (TCGA abbreviation: KIRC), we used Welch’s *t*-test quartile.

To identify motifs enriched and depleted in the rRE catalog, we followed the same method used in the Motif-Scan python module (v1.3.0)^58^. We compared our rRE catalog to Simple Repeats (Tandem Repeat Finder, TRF) as a control. For each unique motif present, we built a contingency table specifying the count of rREs and Simple Repeats with and without the motif. Two one-tailed Fisher’s exact tests were applied to the table to test for significance in both directions, enrichment and depletion. The “stats” module in the Scipy python package (v1.7.0) was used to conduct the significance test. Since multiple hypothesis tests were performed, we applied FDR correction (Benjamini-Hochberg) for multiple hypothesis testing to the p-values, with a cutoff (FDR) of 0.01.

For the comparison of SNVs in COSMIC genes to rREs, we first divided cancer genomes into two categories: rRE cohort and non-rRE cohort. The rRE cohort contains all of the genomes that have at least one rRE detected (n = 615) and the non-rRE cohort contains all of the genomes that have no rREs detected (n = 1897). We then looked at the number of donors in the rRE cohort that have at least one mutation on a given gene (COSMIC Tier 1 genes) i and the number of donors in the non-rRE cohort that have at least one mutation on a given gene i with a contingency table. We then calculated the p-value (Fisher’s exact test) for the significance of associating genes to either rRE or non-rRE cohort. This p-value calculation is repeated for all COSMIC genes and then an FDR at 0.05 significance level (Benjamini-Hochberg) was employed to correct for multiple hypothesis testing.

### Estimation of expansions in the general population

To estimate the frequency of the rREs in the general population, ExpansionHunter Denovo (version 0.9.0) was run on 1000 Genomes Project samples^59^ (n = 2,504) (GRCh38) and Medical Genome Reference Bank^60^ samples (n = 4,010) (GRCh37 lifted over to GRCh38).

The genomic coordinates of the 160 rREs (GRCh37) were padded with 1,000 bp and translated to GRCh38 coordinates with the UCSC LiftOver. Then, these rRE coordinates (GRCh38) were overlapped with loci from the population samples containing the Anchored IRR calls. rREs that overlapped with matching motifs in the population samples were selected for further analysis. We next sought to identify expanded rREs in the population samples to quantify their prevalence. To do so, we converted their global-normalized Anchored IRR values to be comparable to ICGC values. This step was necessary because sequencing read lengths from the PCAWG dataset are generally 100 bp while the read lengths from 1000Genomes and Medical Genome Reference Bank are 150 bp. The conversion follows the formula (Anchored IRR, 100 bp) = 0.5 + 1.5 * (Anchored IRR, 150 bp)^15^. A sample in the population samples was counted as expanded if its Anchored IRR value was greater than the 99th percentile of Anchored IRR values in the normal samples from the PCAWG dataset, a threshold that is comparable to the threshold used to call expansions in tumor samples (**Extended Data Figure 4**). In future rRE catalogs, for the rare instance where the estimated frequency of repeat expansions in the population samples is higher than expected, these data could be used to further filter rREs to improve the detection of cancer-specific repeat expansions.

To compare the length of TRs in normal samples with and without a matching rRE in a tumor sample, donors in the Prost-AdenoCA and Kidney-RCC cohorts whose data are available for download through the Cancer Collaboratory were included (n=253). We used ExpansionHunter (v5.0.0) with the default options to genotype prostate and kidney cancer rREs in the normal samples of the selected donors. When there were two alleles of an rRE in a sample, both alleles were included and treated as distinct data points. For each rRE, we tested whether the distribution of genotypes from donors who have an expansion in their tumor samples differed from donors who did not have an expansion. Student’s t-test was used to compute p-values, and FDR-correction (Benjamini-Hochberg) to adjust for multiple hypothesis testing.

### Association of rREs with gene expression

Matching RNA-seq and WGS data were available for Kidney-RCC, Ovary-AdenoCA, Panc-AdenoCA, and Panc-Endocrine. RNA-seq data from these samples were obtained from DCC (https://dcc.icgc.org/) and values were converted to transcripts per million (TPM). Normalized gene expression (TPM) values were compared for samples with and without an rRE (Welch’s *t*-test, with FDR correction). For isoform analysis, normalized gene expression counts were compared for samples with and without a repeat expansion using the DESeq2 (v1.32.0) package in R v4.0.5. We used the DESeq function to calculate the log_2_ fold change values for 3 isoforms of the *UGT2B7* gene (ENST00000305231.7, ENST00000508661.1, ENST00000502942.1) and performed a Wald test with FDR correction using the Benjamini-Hochberg procedure (threshold *q*-value < 0.01).

### Design, synthesis, and characterization of Syn-TEFs and PAs

Synthetic transcription elongation factors (Syn-TEFs) and polyamides (PAs) were designed to target a GAAA repeat (Syn-TEF3 and PA3) or a control G**G**AA repeat (Syn-TEF4 and PA4). Syn-TEF3, Syn-TEF4, PA3, and PA4 were synthesized and purified to a minimum of 95% compound purity by WuXi Apptec and used without further characterization. HPLC conditions for chemical characterization: 1.0 mL/min, Solvent A: 0.1% (v/v) trifluoroacetic acid (TFA) in H2O, Solvent B: 0.075% (v/v) TFA in acetonitrile, Gemini, Column: C18 5 µm 110A 150*4.6mm. Full results of characterization can be found in **Fig. S2**.

### Treatment of RCC cell lines with synthetic transcription elongation factors (Syn-TEFs)

Caki-1, and 786-O, and Caki-2 cells were obtained from ATCC and grown in RPMI 1640 media with L-glutamine (Gibco Catalog 11875093), supplemented with 10% (v/v) FBS. A498 and ACHN cells were obtained from ATCC and grown in DMEM media with glucose, L-glutamine, and sodium pyruvate (Corning Catalog 10-013-CV), supplemented with 10% (v/v) FBS. RCC-4 cells were obtained from Amato Giacca (Stanford University) and grown in DMEM media with glucose, L-glutamine, and sodium pyruvate (Corning Catalog 10-013-CV), supplemented with 10% (v/v) FBS. Cell lines were confirmed by STR profiling (Genetic Resource Core Facility, Johns Hopkins University) and tested negative for mycoplasma. Cells were seeded in 96-well plates on day 0. On day 1, cells were treated with the indicated molecules. Molecules were dissolved in DMSO (vehicle) and added to cells (0.1% (v/v) DMSO final concentration). On day 4 (72 h later), relative metabolic activity was measured as a proxy for relative cell density, with the Cell Counting Kit (CCK-8; Dojindo Molecular Technologies) per the manufacturer’s instructions. Absorbance (450 nm) of cells treated with molecules was normalized to DMSO (0.1%(v/v)) or no treatment. Absorbance was measured with an Infinite M1000 microplate reader (Tecan, Mannedorf, Switzerland).

For microscopy, Caki-1 and 786-O cells were plated on glass-bottom 96-well plates under standard culture conditions. One day after plating, media containing either no drug, 0.1%(v/v) DMSO, 50 µM Syn-TEF3, or 50 µM Syn-TEF4 was added, and the cells were incubated for 72 hours at 37°C. As a control, wells that received no treatment were incubated with 70%(v/v) ethanol for 30 seconds prior to staining. Cells were then stained with propidium iodide and Hoechst 33342 from the Live-Dead Cell Viability Assay Kit (Millipore Sigma, Catalog CBA415) according to manufacturers’ instructions and immediately imaged at 10x magnification with a 0.17 numerical aperture CFI60 objective on a Keyence BZ-X710 microscope. Four replicates were measured for each treatment condition, and the experiment was repeated three times. Quantitation was conducted using FIJI software. For statistical analyses, a one-way ANOVA with multiple comparisons was conducted with GraphPad Prism.

## Data availability

Whole-genome sequencing data (both short- and long-read DNA sequencing) from 786-O and Caki-1 cell lines are deposited in NCBI with accession PRJNA868795.

## Acknowledgments

This work was supported by NIH grants U2CCA233311 (to M.P.S.) and K99HG011467 (to G.S.E.). G.S.E. was also supported by a Stanford Cancer Institute Postdoctoral Fellowship from the Ellie Guardino Research Fund. Computational support from the Cancer Genomics Cloud (to G.G. and G.S.E.) and an AWS Cloud Research Grant (to G.S.E.). We thank Chiara Sabatti for advice on statistical analysis, Sean O’Connor for preliminary help with data processing, Kevin Van Bortle for advice, and Laura Vanderploeg and Meara Algama for figures. G.S.E. thanks Peter S. Kim for early advice and encouragement.

## Author Contributions

G.S.E. conceived the study. G.S.E., G.G., A.C.F., J.T.L., M.A.E., M.P.S., and M.G. supervised research. G.S.E., G.G., R.A., A.S, E.D., J.P., C.M.B., K.Z., R.K.C.Y., and A.A.E. analyzed data. G.S.E., C.R.H., L.R., A.A., A.A., K.V.K., R.A.K., D.A.S., S.M.W., and T.J.M. conducted wet lab experiments. G.S.E. and M.P.S. wrote the manuscript with input from all the authors.

## Additional Information

**Supplementary Information** is available for this paper.

