## Extended Data Figures 1-10 for "A genome-wide atlas of recurrent repeat expansions in human cancer"

**Extended Data Figure 1. Overview of PCAWG data and analysis with ExpansionHunter De Novo.** a) Distribution of cancer genomes analyzed across 29 human cancers in the PCAWG data. b) Distribution of p-values following candidate recurrent repeat expansion (rRE) analysis with ExpansionHunter Denovo (one-sided Wilcoxon rank-sum test).

a

b

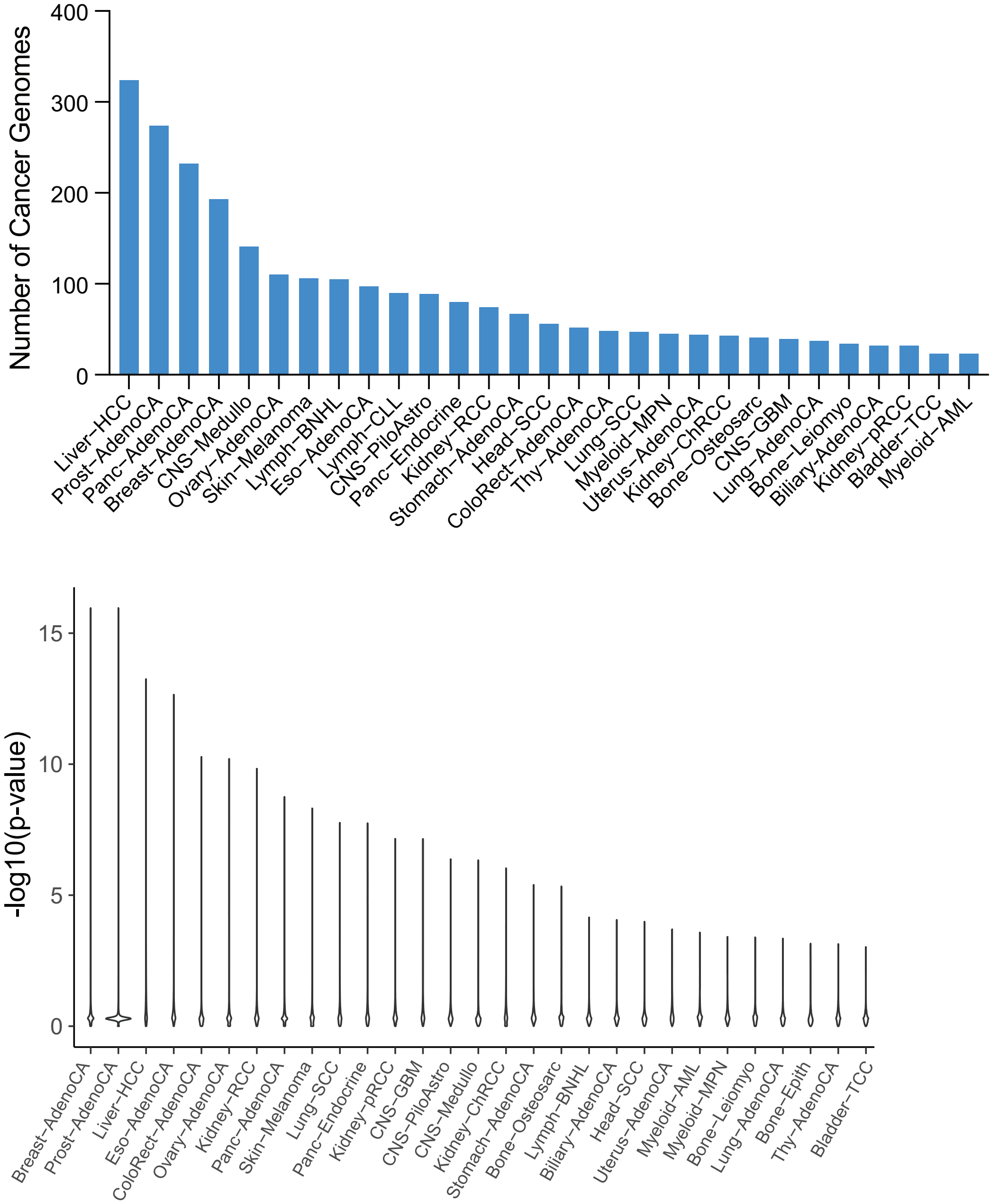

**Extended Data Figure 2. Benchmarking EHdn.** a) Comparison of anchored in-repeat reads (IRRs) to long-read sequencing reads. Long-read sequencing confirmation rate across all tandem repeats (TRs, motifs 2–20 bp), short TRs with motifs from 2–6 bp, and variable number TRs with motifs from 7–20 bp. b) Confirmation rate versus number of anchored IRRs. c) Effect of downsampling on the identification of the rRE in the intron of *UGT2B7* in kidney cancer. Tumor genomes from the PCAWG dataset were downsampled to the specified number. ExpansionHunter De Novo was run, and the resulting Bonferroni-correct p-value is depicted for the given sequencing coverage. d) Estimation of the frequency of repeat expansions in rRE loci in the general population. The number of rREs (count) corresponding to each bin is plotted on the y-axis. Results are from analysis of 1000 Genomes Project samples[33] (n = 2,504) (GRCh38) and Medical Genome Reference Bank[34] samples (n = 4,010).

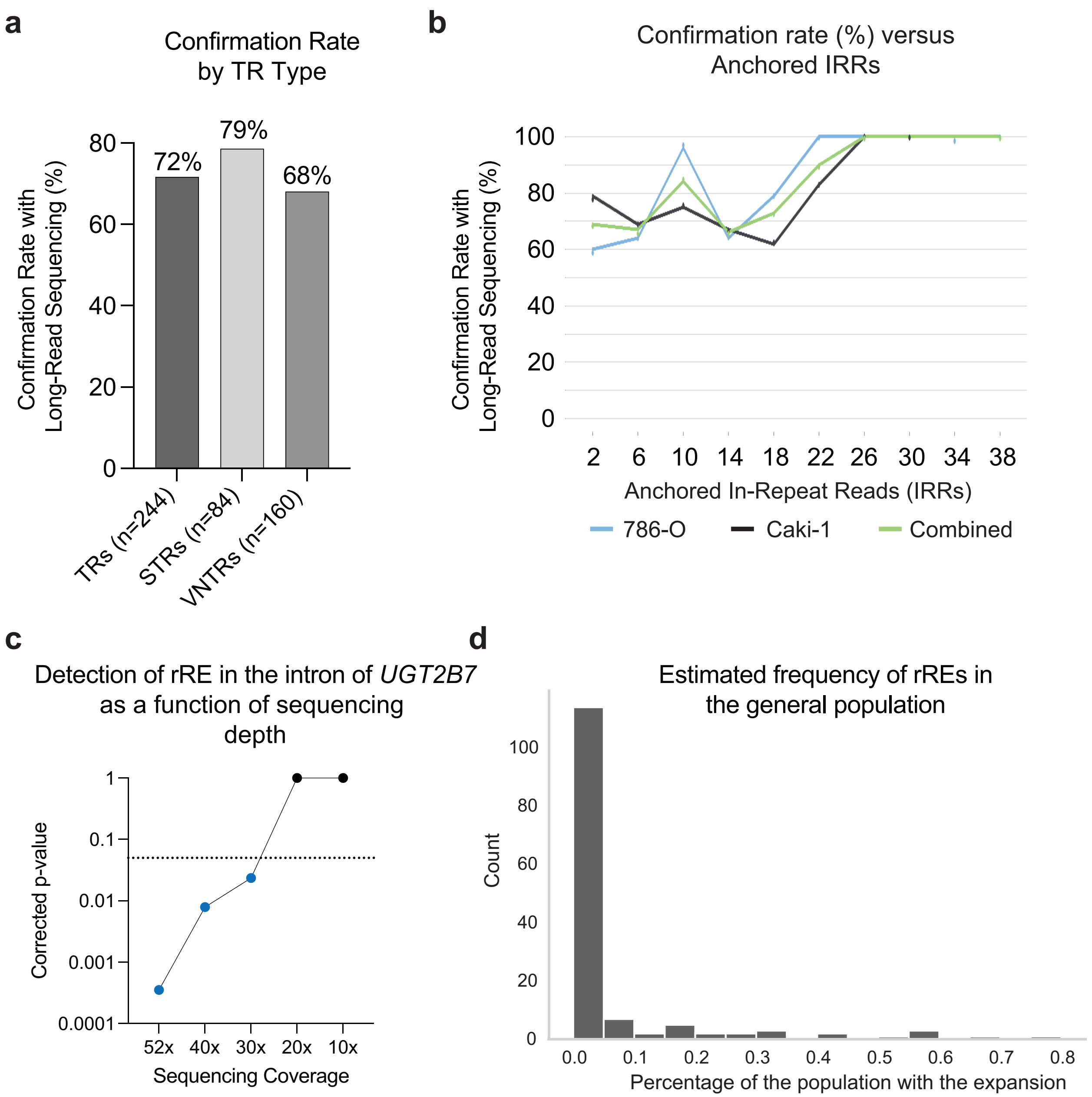
a
Confirmation Rate
by TR Type
Confirmation Rate with
Long-Read Sequencing (%)
72%
79%
68%
80
60
40
20
0
TRs (n=244)
STRs (n=84)
VNTRs (n=160)
b
Confirmation rate (%) versus
Anchored IRRs
Confirmation Rate with
Long-Read Sequencing (%)
100
80
60
40
20
0
2
6
10
14
18
22
26
30
34
38
Anchored In-Repeat Reads (IRRs)
786-O
Caki-1
Combined
c
Detection of rRE in the intron of UGT2B7
as a function of sequencing
depth
Corrected p-value
1
0.1
0.01
0.001
0.0001
52x
40x
30x
20x
10x
Sequencing Coverage
d
Estimated frequency of rREs in
the general population
Count
100
80
60
40
20
0
0.0
0.1
0.2
0.3
0.4
0.5
0.6
0.7
0.8
Percentage of the population with the expansion

**Extended Data Figure 3. Local read depth normalization of recurrent repeat expansion (rRE) candidates.** a) Examples of read depth before and after local normalization. b) Examples of anchored in-repeat read (IRRs) before and after local normalization. c) Workflow to identify rREs. d) Detection rate in an independent cohort of samples.

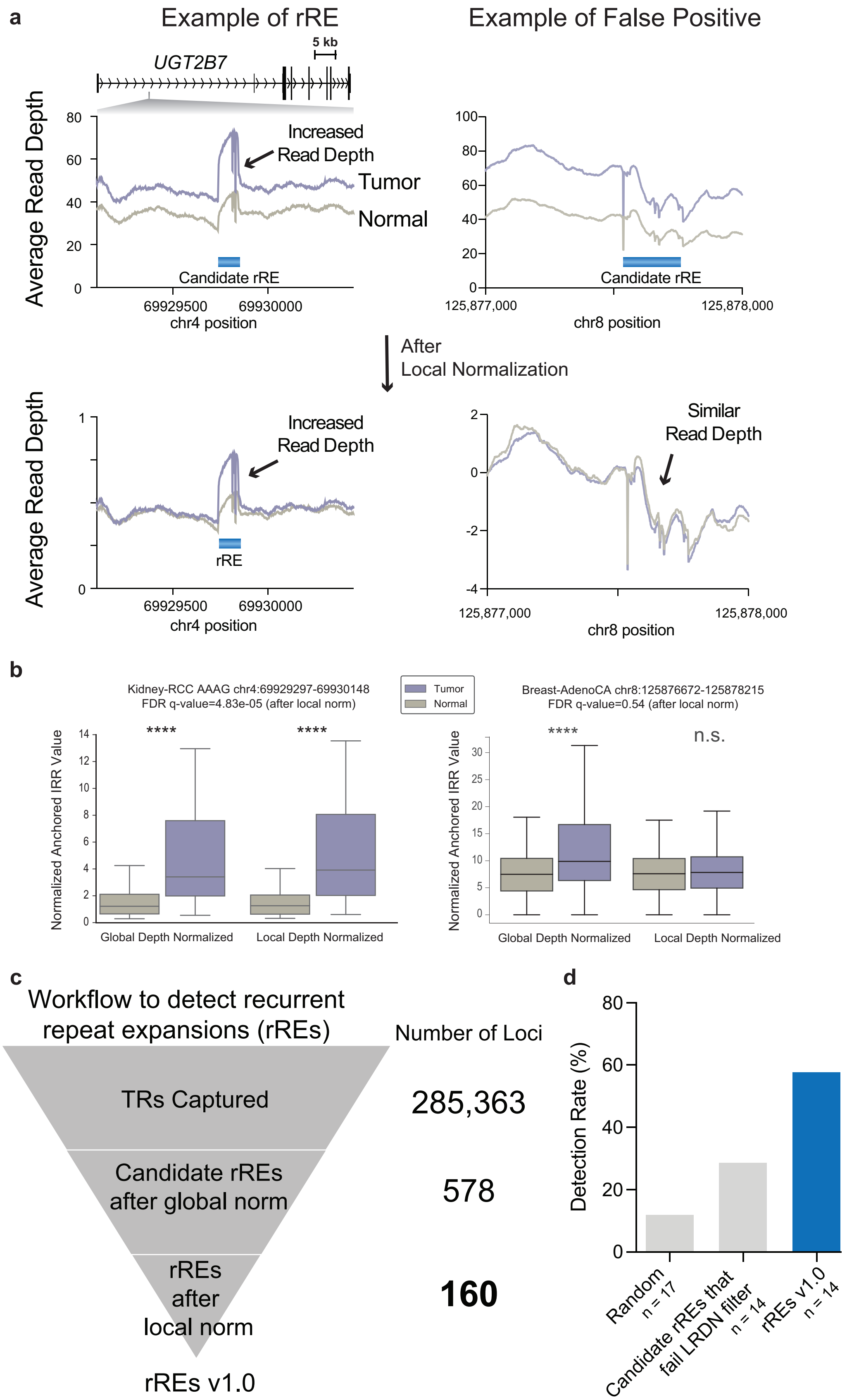
a
Example of rRE
Example of False Positive
5 kb
UGT2B7
Average Read Depth
Increased
Read Depth
Tumor
Normal
Candidate rRE
69929500
69930000
chr4 position
125,877,000
125,878,000
chr8 position
After
Local Normalization
rRE
Similar
Read Depth
b
Kidney-RCC AAAG chr4:69929297-69930148
FDR q-value=4.83e-05 (after local norm)
Tumor
Normal
Breast-AdenoCA chr8:125876672-125878215
FDR q-value=0.54 (after local norm)
Normalized Anchored IRR Value
****
n.s.
Global Depth Normalized
Local Depth Normalized
c
Workflow to detect recurrent
repeat expansions (rREs)
Number of Loci
TRs Captured
285,363
Candidate rREs
after global norm
578
rREs
after
local norm
160
rREs v1.0
d
Detection Rate (%)
Random
n = 17
Candidate rREs that
fail LRDN filter
n = 14
rREs v1.0
n = 14

**Extended Data Figure 4. Benchmarking LRDN and EHdn.** a) Benchmarking the local read depth normalization filter. b) The Anchored IRR quotient was calculated as (Tumor Anchored IRR – Normal Anchored IRR)/(Normal Anchored IRR + 1). Dashed line at 2.5 indicates the threshold for calling a locus as a repeat expansion in a cancer genome. c) ExpansionHunter was used to estimate repeat sizes from short-read sequencing data, and the results were visualized with REViewer (see **Methods**). The allele with the longest repeat tract for normal and tumor samples is shown. The TR is depicted in red, and the flanking regions are depicted in blue.

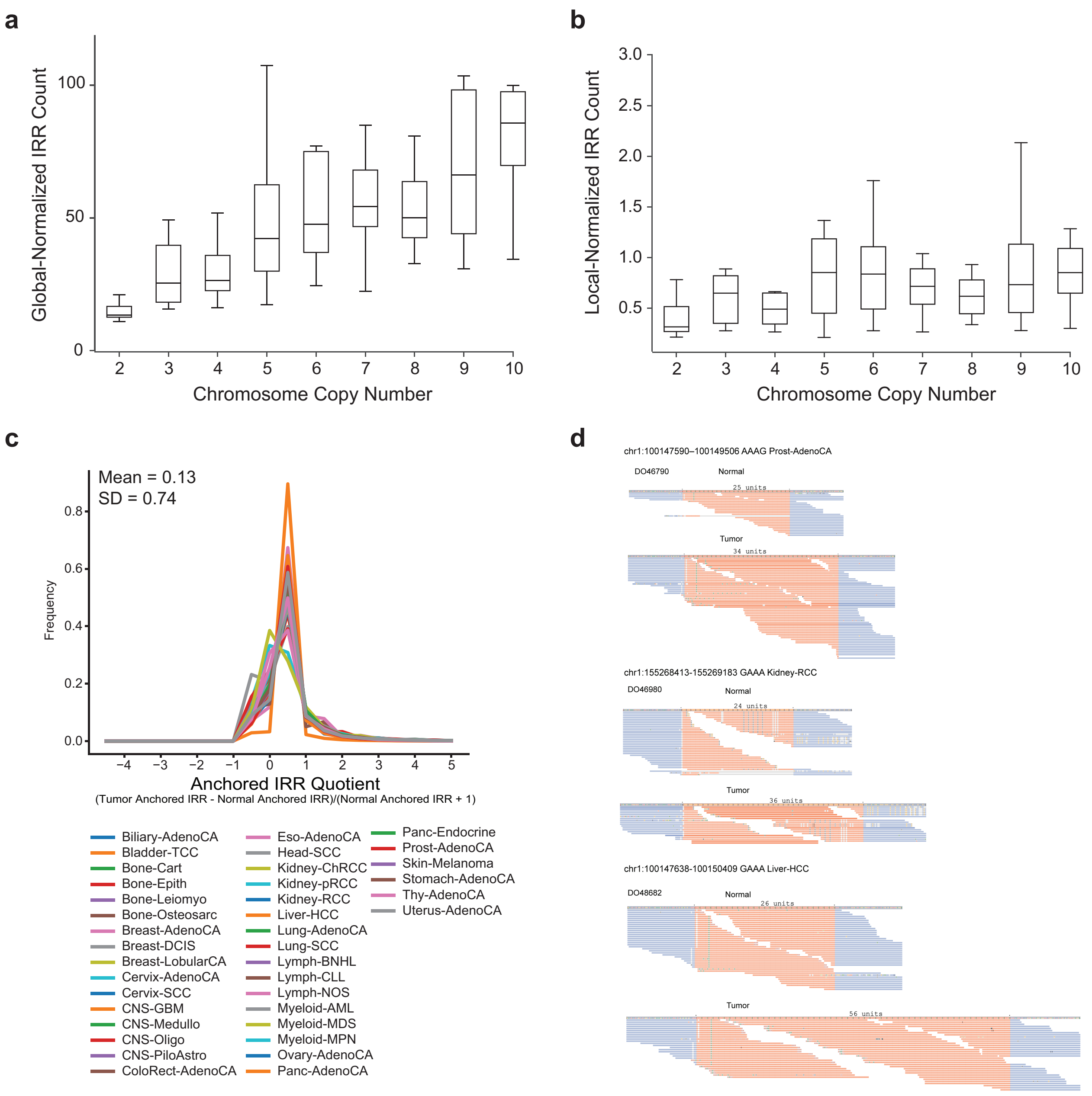
a
Global-Normalized IRR Count
0
50
100
2
3
4
5
6
7
8
9
10
Chromosome Copy Number
b
Local-Normalized IRR Count
0.5
1.0
1.5
2.0
2.5
3.0
Chromosome Copy Number
c
Mean = 0.13
SD = 0.74
Frequency
0.0
0.2
0.4
0.6
0.8
−4
−3
−2
−1
0
1
2
3
4
5
Anchored IRR Quotient
(Tumor Anchored IRR - Normal Anchored IRR)/(Normal Anchored IRR + 1)
Biliary-AdenoCA
Bladder-TCC
Bone-Cart
Bone-Epith
Bone-Leiomyo
Bone-Osteosarc
Breast-AdenoCA
Breast-DCIS
Breast-LobularCA
Cervix-AdenoCA
Cervix-SCC
CNS-GBM
CNS-Medullo
CNS-Oligo
CNS-PiloAstro
ColoRect-AdenoCA
Eso-AdenoCA
Head-SCC
Kidney-ChRCC
Kidney-pRCC
Kidney-RCC
Liver-HCC
Lung-AdenoCA
Lung-SCC
Lymph-BNHL
Lymph-CLL
Lymph-NOS
Myeloid-AML
Myeloid-MDS
Myeloid-MPN
Ovary-AdenoCA
Panc-AdenoCA
Panc-Endocrine
Prost-AdenoCA
Skin-Melanoma
Stomach-AdenoCA
Thy-AdenoCA
Uterus-AdenoCA
d
chr1:100147590–100149506 AAAG Prost-AdenoCA
DO46790
Normal
25 units
Tumor
34 units
chr1:155268413-155269183 GAAA Kidney-RCC
DO46980
Normal
24 units
Tumor
36 units
chr1:100147638-100150409 GAAA Liver-HCC
DO48682
Normal
26 units
Tumor
56 units

**Extended Data Figure 5**. **Association of rREs with genetic features.** a) Correlation of rREs with MSI-High cancers. b and c) Association of rREs with Mutational Signatures. b) Correlation between DBS2 and the number of rREs detected. c) Correlation between DBS2 and the number of rREs detected when Lung-SCC data are omitted from the analysis.

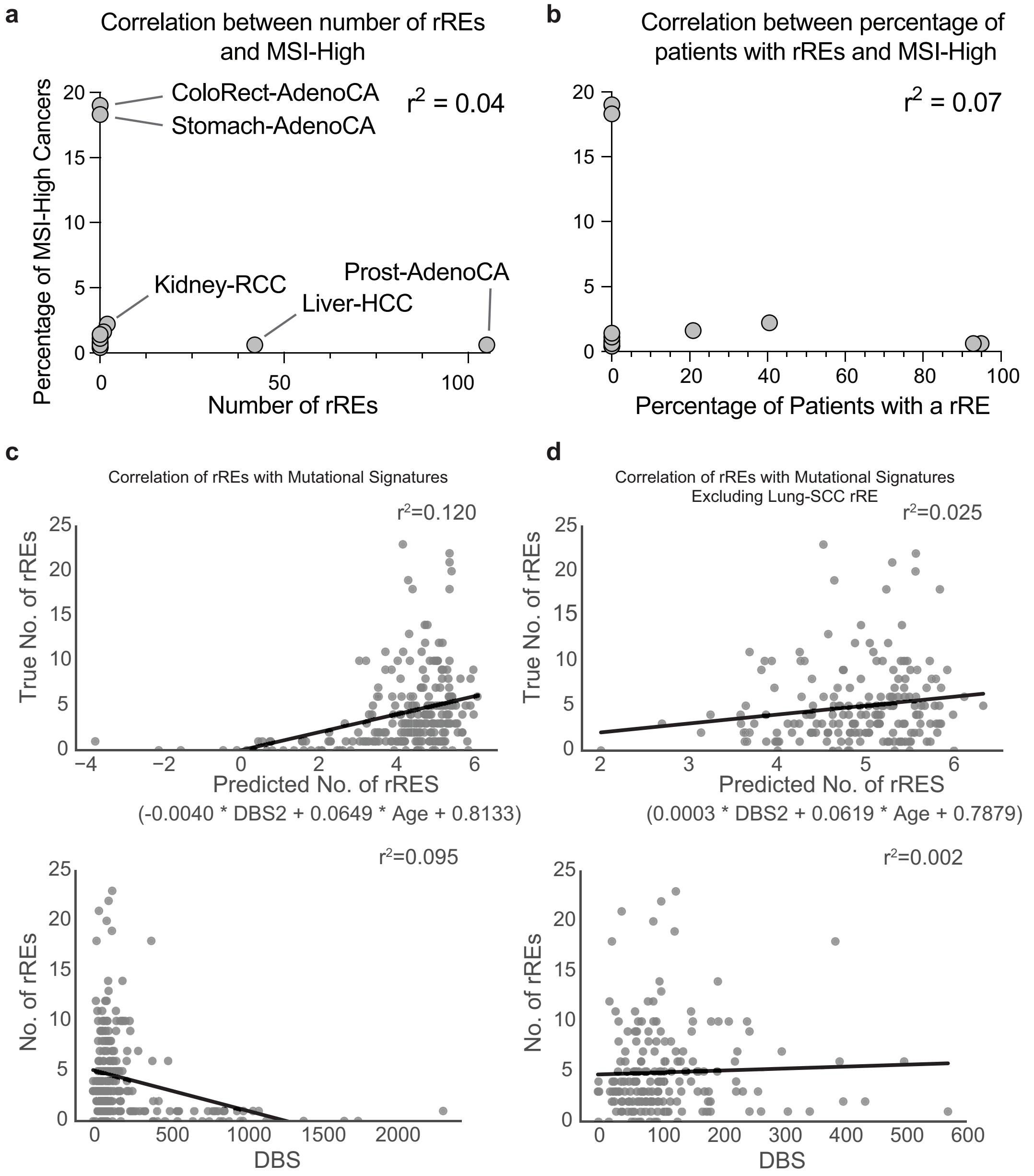

**Extended Data Figure 6. Distribution of rREs across the genome.** a) Distance of rREs to the nearest centromere or telomere. b) Distribution of rREs across early- and late-replicating regions of the genome. Welch's *t*-test (not significant). c) Circos plot depicting (from outside to inside) p-value of rREs, location of rREs where darker shading indicates the rRE observed across 3 cancers, early and late replicating regions (yellow and purple, respectively), and simple sequence repeats. This plot depicts the overlay between different data types and the distribution of rREs across the genome.

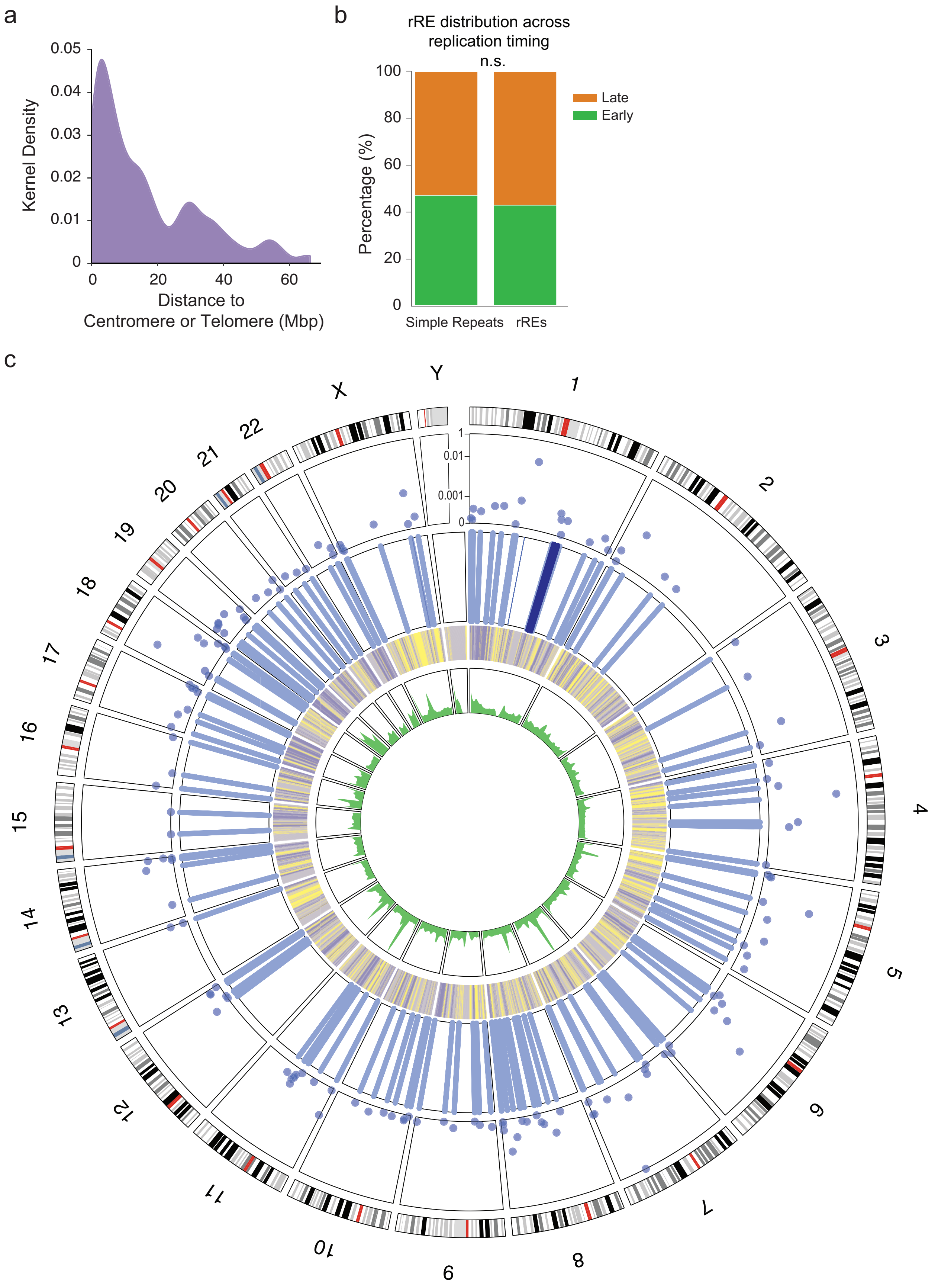
a
0.05
0.04
0.03
0.02
0.01
0
Kernel Density
0
20
40
60
Distance to
Centromere or Telomere (Mbp)
b
rRE distribution across
replication timing
n.s.
100
80
60
40
20
0
Percentage (%)
Late
Early
Simple Repeats
rREs
c
Y
X
22
21
20
19
18
17
16
15
14
13
12
11
10
9
8
7
6
5
4
3
2
1
0.01
0.001
0

**Extended Data Figure 7. Molecular Features of rREs**. a) Overlap of rREs with other datasets. The fraction of rREs overlapping with other catalogs of TRs and genomic instability. From left to right in the figure, recurrently altered STRs in cancer (Supplementary Data 14 from Cortes-Ciriano et al, & Park, PMID: 28585546), extrachromosomal circular DNA (ecDNA, circular amplification events from Supplementary Table 1, Wu et al, & Mischel, PMID: 31748743), unstable STRs in cancer (Supplementary Table 10 from Hause et al, & Salipante, PMID: 27694933), eSTRs (Supplementary Data 1, Fotsing et al, & Gymrek, PMID: 31676866), and microDNA (From C4-2, ES2, LNCaP, OVCAR8, and PC-3 cells, Dillon et al, & Dutta, PMID: 26051933). The PubMed ID for each corresponding manuscript is included in the figure. For the overlap of rREs with microDNA, we looked at loci that we attempted to detect in an independent cohort of cancer samples, and we found that we tested 11 loci. Of the 11 rREs tested, 8 (72%) were detected in the independent cohort of cancer samples. b) Distribution of rRE motif length across cancer types. b and c) Association of rREs with regulatory elements. b) Distance of simple sequence repeats and rREs to the nearest candidate cis-regulatory elements (cCREs). Key: promoter-like signature (P), proximal enhancer-like signature (p), distal enhancer-like signature (d), DNase-H3K4me3 (D), and CTCF-only (C). c) Signal tracks depicting rREs near regulatory elements. d) Association between rREs in prostate cancer and risk loci in prostate cancer. Signal trace showing a prostate cancer rRE and a risk locus.

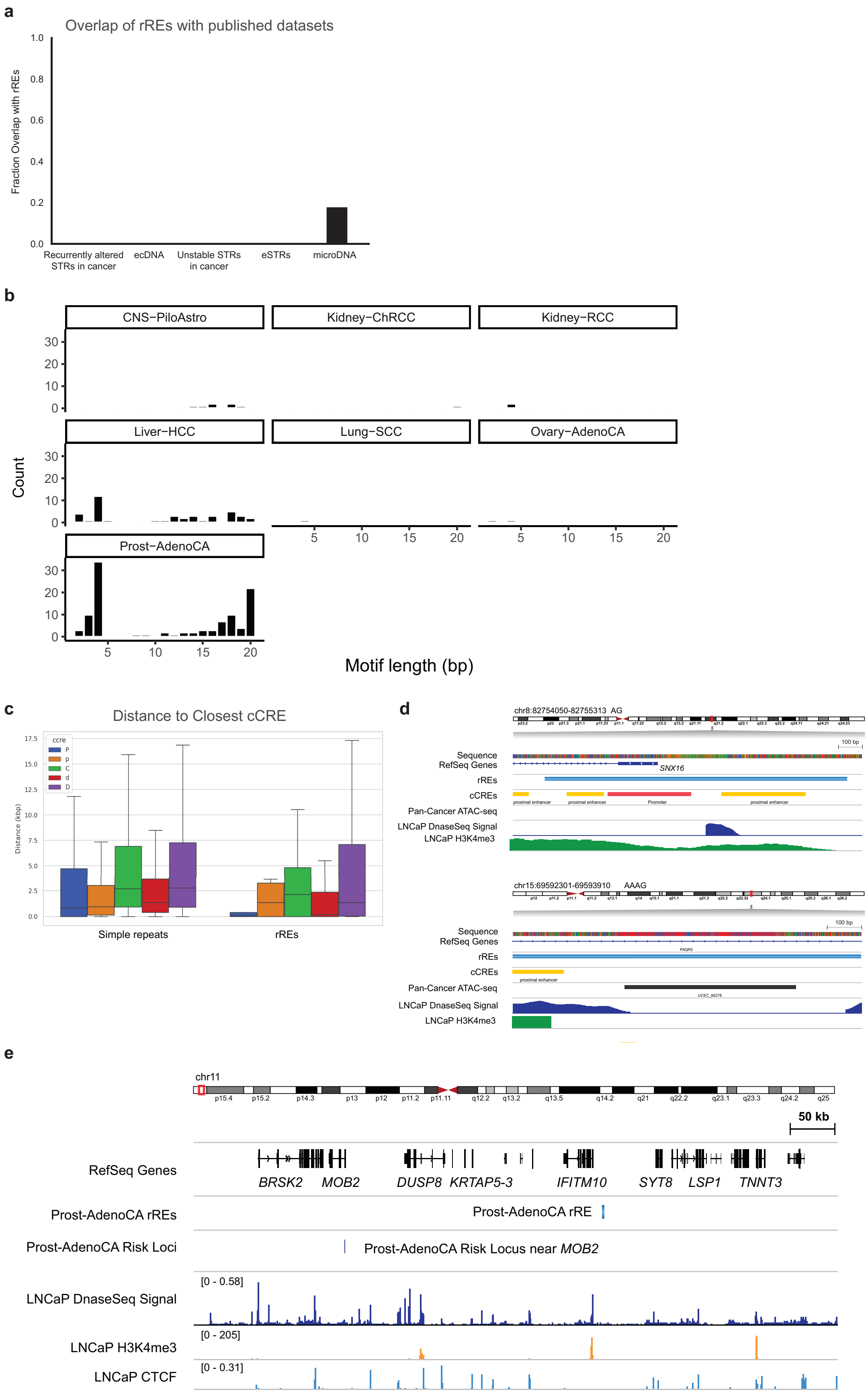
a
Overlap of rREs with published datasets
Fraction Overlap with rREs
1.0
0.8
0.6
0.4
0.2
0.0
Recurrently altered STRs in cancer
ecDNA
Unstable STRs in cancer
eSTRs
microDNA
b
CNS−PiloAstro
Kidney−ChRCC
Kidney−RCC
Liver−HCC
Lung−SCC
Ovary−AdenoCA
Prost−AdenoCA
Count
Motif length (bp)
c
Distance to Closest cCRE
Distance (kbp)
ccre
P
p
C
d
D
Simple repeats
rREs
d
chr8:82754050-82755313 AG
Sequence
RefSeq Genes
SNX16
rREs
cCREs
proximal enhancer
Promoter
Pan-Cancer ATAC-seq
LNCaP DnaseSeq Signal
LNCaP H3K4me3
chr15:69592301-69593910 AAAG
e
chr11
50 kb
RefSeq Genes
BRSK2
MOB2
DUSP8
KRTAP5-3
IFITM10
SYT8
LSP1
TNNT3
Prost-AdenoCA rREs
Prost-AdenoCA rRE
Prost-AdenoCA Risk Loci
Prost-AdenoCA Risk Locus near MOB2
LNCaP DnaseSeq Signal
[0 - 0.58]
LNCaP H3K4me3
[0 - 205]
LNCaP CTCF
[0 - 0.31]

**Extended Data Figure 8.** a) Analysis of *UGT2B7* GAAA rRE in patients with clear cell RCC. N, Normal tissue; T, Tumor tissue. For gel source data, see **Fig. S1**. b) *UGT2B7* in RCC patients. b) Expression of *UGT2B7* (transcripts per million, TPM) in RCC samples as a function of the detection of the rRE in *UGT2B7*. P value computed with Welch's *t*-test. c) Kaplan-Meier survival plots of RCC patients stratified by rRE in the intron of *UGT2B7*. P value computed with Welch's *t*-test.

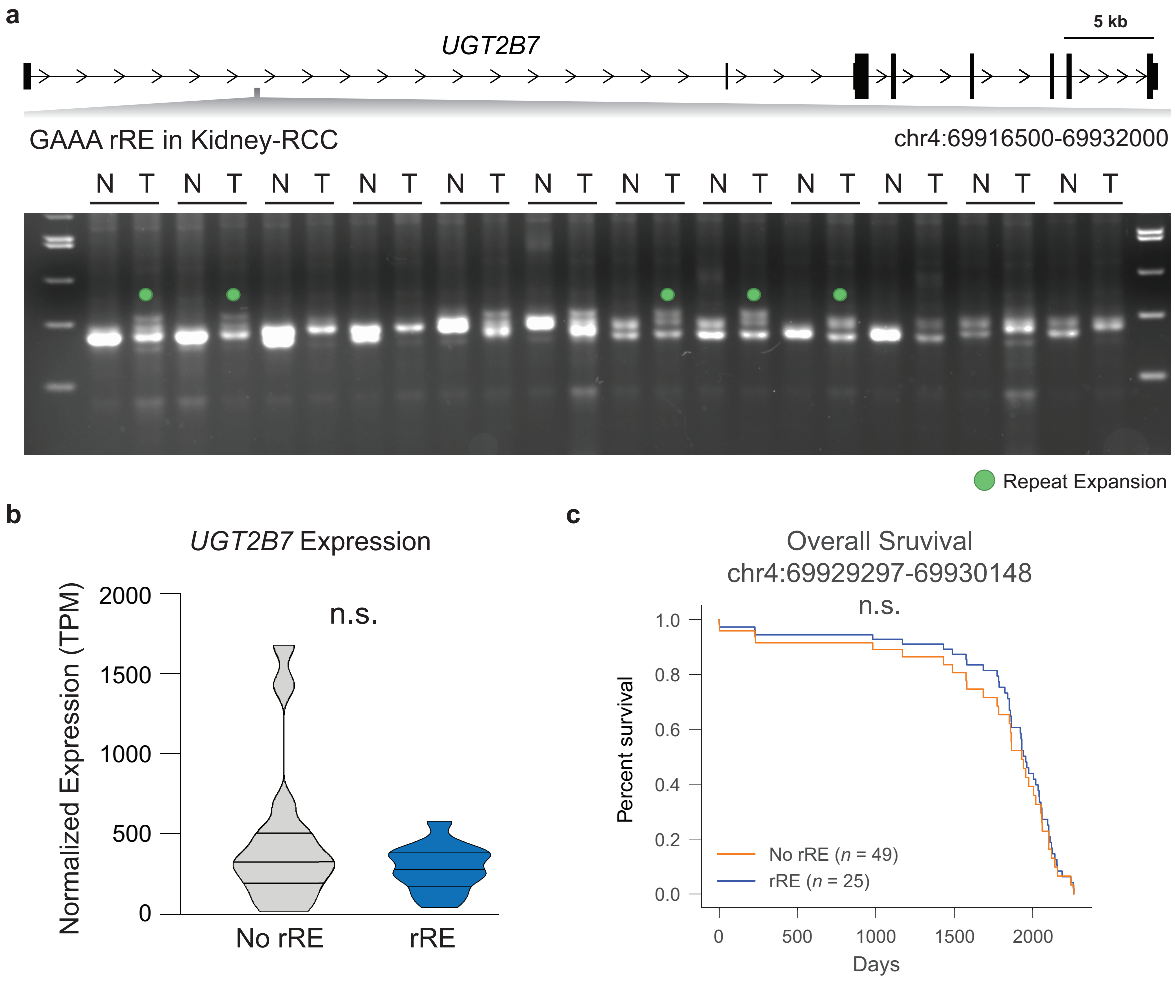

**Extended Data Figure 9. Association of rREs with cytotoxic activity.** P values computed with Welch's *t*-test with FDR correction (Benjamini-Hochberg).

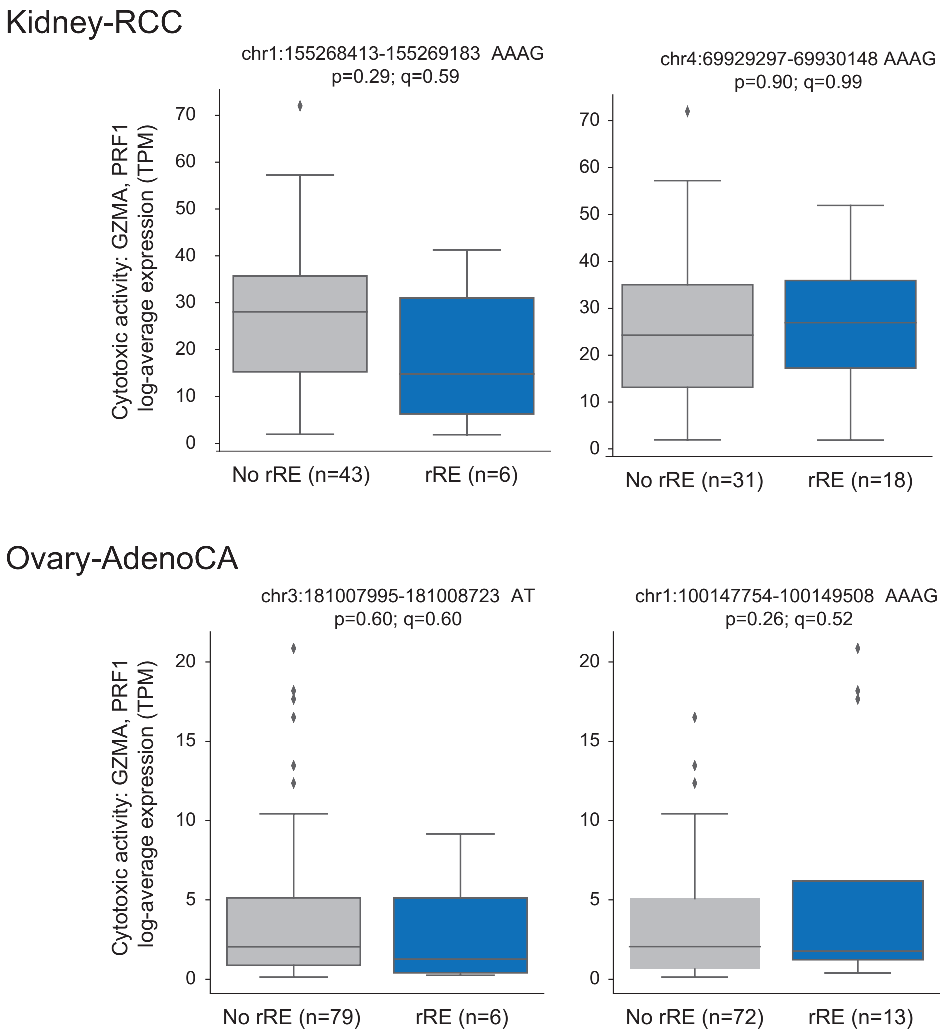

**Extended Data Figure 10. Syn-TEF treatment of RCC cell lines.** a) Relative cell density of RCC cell lines following treatment (72 h) with compounds (50 µM Syn-TEF or 0.1% DMSO vehicle, as indicated). Results are mean ± SEM ($n$ = 4). b) Quantitation of the percentage of propidium iodide-positive cells. Whiskers represent minimum and maximum values. * $p < 0.05$, *** $p < 0.0001$. P values were calculated from a one-way ANOVA with multiple comparisons. c) Live cell microscopy of Caki-1 and 786-O cells stained with propidium iodide (red) and Hoechst 33342 (blue). Scale bars, 100 µm.

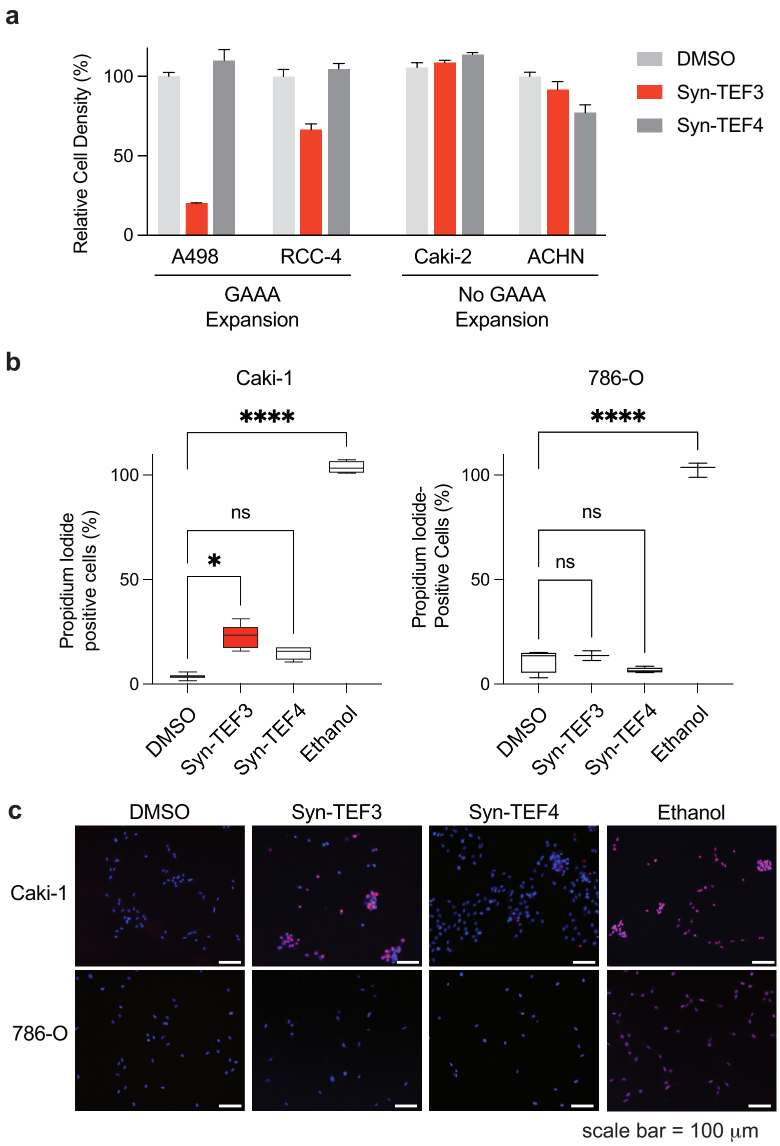
